# *MYC* dosage compensation is mediated by miRNA-transcription factor interactions in aneuploid cancer

**DOI:** 10.1101/2021.04.20.440572

**Authors:** ManSai Acón, Carsten Geiß, Jorge Torres-Calvo, Diana Bravo-Estupiñan, Guillermo Oviedo, Jorge L Arias-Arias, Luis A Rojas-Matey, Edwin Baez, Gloriana Vásquez-Vargas, Yendry Oses-Vargas, José Guevara-Coto, Andrés Segura-Castillo, Francisco Siles-Canales, Steve Quirós-Barrantes, Anne Régnier-Vigouroux, Pedro Mendes, Rodrigo Mora-Rodríguez

## Abstract

We hypothesize that dosage compensation of critical genes arises from systems-level properties for cancer cells to withstand the negative effects of aneuploidy. We identified several candidate genes in cancer multi-omics data and developed a biocomputational platform to construct a mathematical model of their interaction network with miRNAs and transcription factors, where the property of dosage compensation emerged for *MYC* and was dependent on the kinetic parameters of its feedback interactions with three micro-RNAs. These circuits were experimentally validated with a novel genetic tug-of-war technique by overexpressing an exogenous *MYC* leading to over-expression of the three microRNAs involved and down-regulation of endogenous *MYC.* In addition, *MYC* overexpression or inhibition of its compensating miRNAs led to dosage-dependent cytotoxicity in *MYC*-amplified colon cancer cells. Finally, we identified negative correlation of *MYC* dosage compensation with patient survival in TCGA breast cancer patients, highlighting the potential of this mechanism to prevent aneuploid cancer progression.

**Highlights:** The systems-level property of gene dosage-compensation emerges *in silico* in miRNA-transcription factor networks depending on the kinetic parameters of its interactions.

We established a criterion to identify compensated candidate genes with low variation in expression despite high copy number variation.

BioNetUCR is a novel biocomputational platform to model miRNA-transcription factor interactions

We present a novel genetic tug-of-war technique to experimentally validate gene dosage compensation at the transcriptional level.

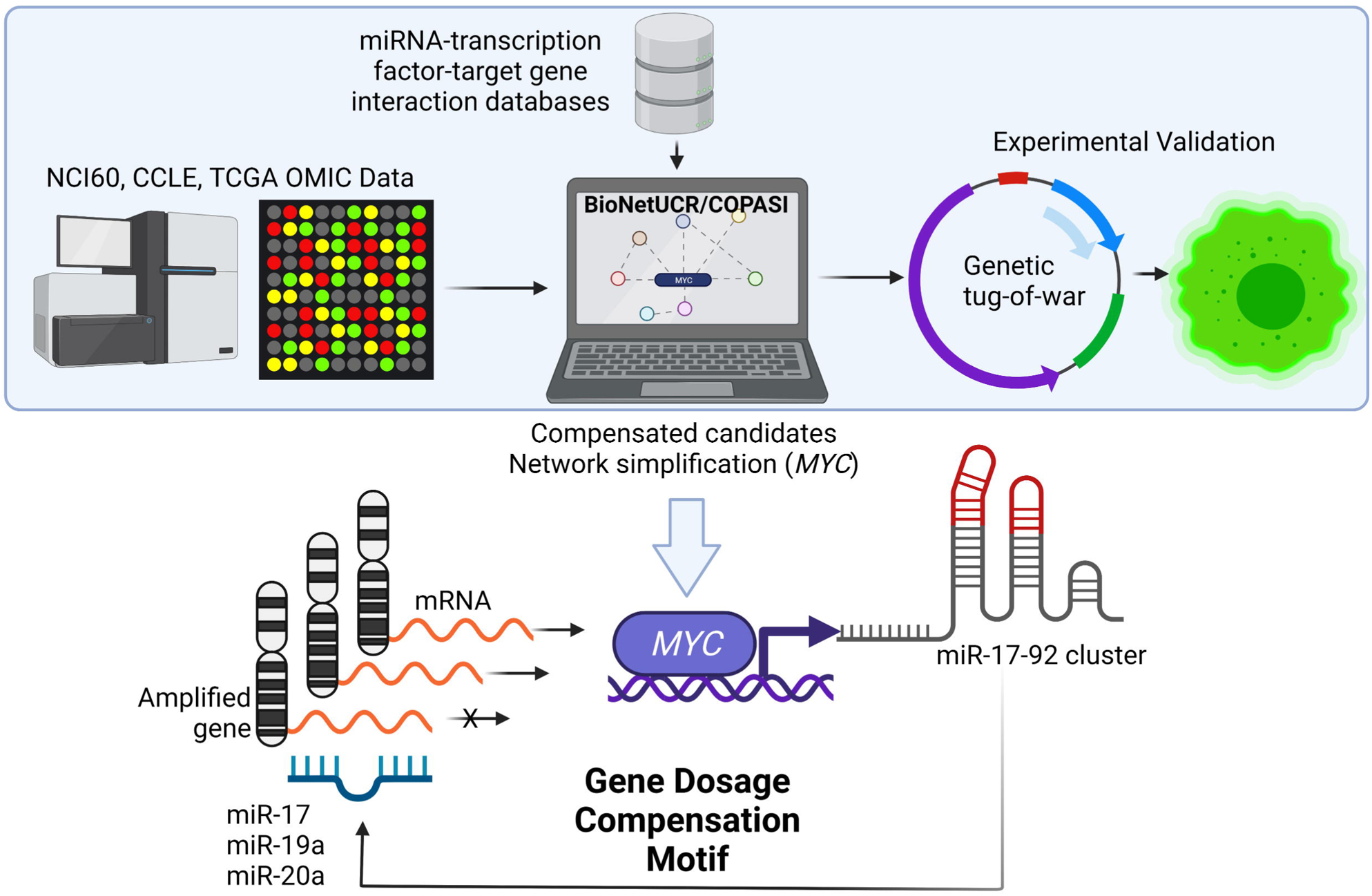

## Introduction

Aneuploidy is a genomic alteration characterized by partial or whole gains and losses of chromosomes (Hanahan and Weinberg, 2011). This results in lethality for normal cells and entire organisms, representing the most frequent cause of miscarriages and intellectual disabilities in humans (Hardy and Hardy, 2015), (Kojima and Cimini, 2019), (Sheltzer and Amon, 2011). Aneuploidy is also the major source of autocatalytic genomic instability, where the cells with the highest aneuploidy have also the highest genomic instability (Duesberg et al., 1998). This leads to many defects at the cellular level such as: decreased cell proliferation and viability, increased proteotoxic stress, metabolic requirements, lactate production and induced recombination defects. The most plausible explanation for all those negative effects of aneuploidy is given by the alteration of the gene dosage, where gains or losses of entire chromosomes alter the dosage of many genes, leading to unbalanced load of critical proteins, altering energetic requirements and protein homeostasis (Sheltzer and Amon, 2011).

Aneuploidy is also a hallmark of cancer. It is present in 90% of solid tumors and 75% of hematologic tumors. Interestingly, aneuploidy was considered by early reports as the driving force of cancer genomic evolution, and seems to have a role in oncogenesis because it appears before malignant transformation, as there are clonal genetic alterations that compromise the fidelity of chromosomal segregation as well as inherited mutations in checkpoints that lead to aneuploidy predisposing to cancer (Sheltzer and Amon, 2011). It is therefore unknown how cancer cells deal with so much aneuploidy whereas normal cells are very sensitive. In fact, aneuploidy autocatalyzes genomic instability of transforming cancer cells leading to cell death due to catastrophic errors (Solé and Deisboeck, 2004). However, in rare occasions, a specific combination of alterations is met, where chromosomes are gained or lost to restore the balance and maintain viability (Ozery-Flato et al., 2011) suggesting the presence of a stable mechanism, whose function has to be maintained to ensure survival.

A hint about the nature of these mechanisms developed by cancer cells to minimize the negative effects of aneuploidy was provided by the fact that hyperdiploidy (higher than normal chromosome number) is more frequently observed than hypodiploidy (lower than normal chromosome number) (Cimini, 2008; Weaver and Cleveland, 2006). Thus, in most cases in which aneuploidy occurs, cells possess a number of chromosomes that is above the norm. A possible explanation is given by the gene dosage compensation hypothesis, a mechanism by which the expression of certain genes is modulated to balance for differences in gene dosage when extra chromosomes are present due to aneuploidy (Kojima and Cimini, 2019).

Gene dosage compensation was described very early for other organisms, as a means to balance the negative effects of aneuploidy (Devlin et al., 1982). Indeed, the concept of gene dosage compensation, or gene dosage balance, is a widespread phenomenon that was discovered in the early days of genetics and there is accumulating evidence that it has an effect on gene expression, quantitative traits, aneuploid syndromes, population dynamics of copy number variants and differential evolutionary fate of genes after partial or whole-genome duplication (for more thorough reviews see (Birchler and Veitia, 2012) (El-Brolosy and Stainier, 2017)).

There are several potential mechanisms of dosage compensation at the many levels of the path that leads from a gene to a final product. At the protein synthesis level, the effects of gene dosage compensation are hypothesized to result from stoichiometric differences among members of macromolecular complexes, the interactome, and signaling pathways (Birchler and Veitia, 2012; Veitia et al., 2008). In this sense, gene dosage compensation represents a compensatory mechanism that may ameliorate the imbalanced gene expression and restore protein homeostasis in aneuploid cells. Indeed, the multiple consequences of aneuploidy are mainly regulated at the protein level due to a side-effect of protein folding defects and increased protein degradation by proteosome and autophagy (Donnelly and Storchová, 2014). An approach to identify genes with dosage compensation by increasing the copy number of individual genes using the genetic tug-of-war technique demonstrated that 10% of the yeast genome have gene dosage compensation at the protein level, and consists predominantly of subunits of multi-protein complexes, which are regulated in a stoichiometry-dependent manner (Ishikawa et al., 2017). In aneuploid cancer, it has been shown that messenger RNA (mRNA) levels generally correlate well with an increased DNA copy number (gene dosage) but these changes are not reflected at the protein level for several genes (Stingele et al., 2012). Indeed, gene dosage compensation in cancer cells has been recently demonstrated at the protein level, whereby protein aggregation mediates the stoichiometry of protein complexes in aneuploid cells. This study shows that excess subunits are either degraded or aggregated, which is nearly as effective as protein degradation at lowering the level of functional proteins (Brennan et al., 2019).

The regulation of gene transcription might be another effective mechanism to compensate the gene dosage changes in aneuploid cells, as it maintains the stoichiometry and preserves the energy required for transcription, translation, and eventual degradation of the extra proteins (Donnelly and Storchová, 2014). At this level, several experimental models have been used to investigate the effects of aneuploidy on gene expression, with contradicting findings. In some of these studies it has been identified that the acquisition of an extra chromosome results in a proportional increase in the expression of genes from that chromosomal unit (Gao et al., 2007; Hughes et al., 2000; Lu et al., 2011; Mao et al., 2003; Pavelka et al., 2010; Pollack et al., 2002; Ried et al., 2012; Stingele et al., 2012; Torres et al., 2007; Tsafrir et al., 2006; Virtaneva et al., 2001; Williams et al., 2008; Xu et al., 2010). In contrast, other studies have reported that certain genes on the aneuploid chromosomes display expression levels similar to those observed in the euploid condition or lower than expected from their dosage (Chikashige et al., 2007; Kahlem et al., 2004; Lyle et al., 2004; Sun et al., 2013a, 2013b; Vacík et al., 2005) Moreover, the degree of compensation may vary and it may occur simultaneously with effects on other chromosomes (for review see (Kojima and Cimini, 2019). Despite those multiple findings, the gene dosage compensation is still a phenomenon in search of mechanisms (El-Brolosy and Stainier, 2017).

Among the group of genomic elements categorized as non-coding RNAs, micro-RNAs (miRNAs) are one of the most predominant represented molecules in this group, and have been widely studied due to their clinical importance. miRNAs are small endogenous RNA molecules that bind mRNAs and repress gene expression (Fabian et al., 2010). A typical miRNA is processed from a long primary RNA sequence to a short mature functional transcript around 22 nucleotides in length. A common characteristic of a miRNA is its ability to pleiotropically target potentially the expression of hundreds or even thousands of genes (Hanna et al., 2019) and their target genes can also be regulated by several miRNAs (Ritchie et al., 2013). Current estimates indicate that the human genome contains 1917 annotated hairpin precursors, and 2654 mature sequences of miRNAs (Kozomara et al., 2019), estimated to directly regulate >60% of human mRNAs (Kim et al., 2016). In consequence, there is a good possibility that miRNA-transcription factor interactions may regulate the expression of genes amplified or deleted in cancer. Therefore, we hypothesize that gene dosage compensation in cancer can be mediated, at least in part, by the emerging properties of complex miRNA-TF networks, controlling the expression of key genes that have altered copy number.

In the present work we studied a potential mechanism of gene dosage compensation in aneuploid tumoral cells with the aid of models constructed by a biocomputational platform. A criterion was established to identify candidate genes with low tolerance in variation of gene expression despite high tolerance in CNV (copy number variations). These genes were input to construct the models to identify compensation circuits mediated by miRNA-TF interactions. We report the transcriptional gene dosage compensation of *MYC*, *STAT3*, *STAT5A* and *STAT5B* mediated *in silico* by an emergent property of feedback and feedforward loops. Thereby, we critically established that the network topology alone is not enough, as the interactions forming these network motifs have a specific set of kinetic parameters enabling compensation, strengthening the need for full dynamic models. We experimentally validated the compensation circuits for *MYC* using a novel genetic tug-of-war approach and determined that this mechanism can be blocked by depleting key miRNAs, affecting in a larger extent those cells with higher copy numbers of those compensated genes. Our analysis of CCLE data confirmed *MYC* dosage compensation across multiple cancer types and the study of TCGA indicated that *MYC* dosage compensation negatively correlates with patient survival.

Thus, we propose that a regulatory network mediated by miRNAs compensates for gene dosage changes in aneuploid cancer cells. We suggest that the manipulation of specific nodes of this miRNA-based regulatory network could block gene dosage compensation, representing a novel type of specific targeting of aneuploid cancer cells.

## Results

### Candidate genes with low tolerance to variation in their RNA expression despite high copy number variation are present across the cancer genomes of the NCI60 panel

The identification of dosage-compensated candidate genes is not trivial. A CNV-buffering score was proposed for aneuploid yeast strains as the dosage-compensated genes had higher variation in copy number but a constrained expression level (Hose et al., 2016). We implemented a similar criterion to identify genes under putative dosage compensation by comparing copy number, gene expression and proteomic data of all genes in the NCI60 panel, a collection of 59 cell lines of 7 types of cancer which are fully characterized at different omic levels (Shoemaker, 2006). We considered input data including high resolution data of Copy Number Variation (CNV) of the NCI60 Cancer Cell lines from 4 different platforms (Bussey et al., 2006), the Gene Transcript (RNA) Average Intensities of 5 Platforms (Gmeiner et al., 2010), and the protein levels (Protein) of a global proteome analysis of the NCI60 cell line panel (Gholami et al., 2013). Figure 1A-left shows the variation of the absolute values of DNA copy number, RNA expression and Protein expression. Next, we calculated the average RNA or protein expression of the diploid cell lines (copy number between −0.25 and 0.25) for each gene to obtain the average diploid expression for every gene. Afterwards, we normalized all expression data against this diploid average expression for each gene and calculated log2 transformed values. Using this approach, we could directly and simultaneously compare the changes in DNA Copy Number, RNA and Protein expression across the NCI60 panel to look for genes with high variations in DNA copy numbers but a low tolerance to variation in RNA or protein expression as they could represent candidate targets under dosage compensation (Figure 1A). We plotted the Standard Deviation (SD) of the DNA, RNA and Protein values across the 59 cell lines and observed that several dots separated from the main gene population as they present high SD of DNA levels (Figure 1B).

**Figure 1.**
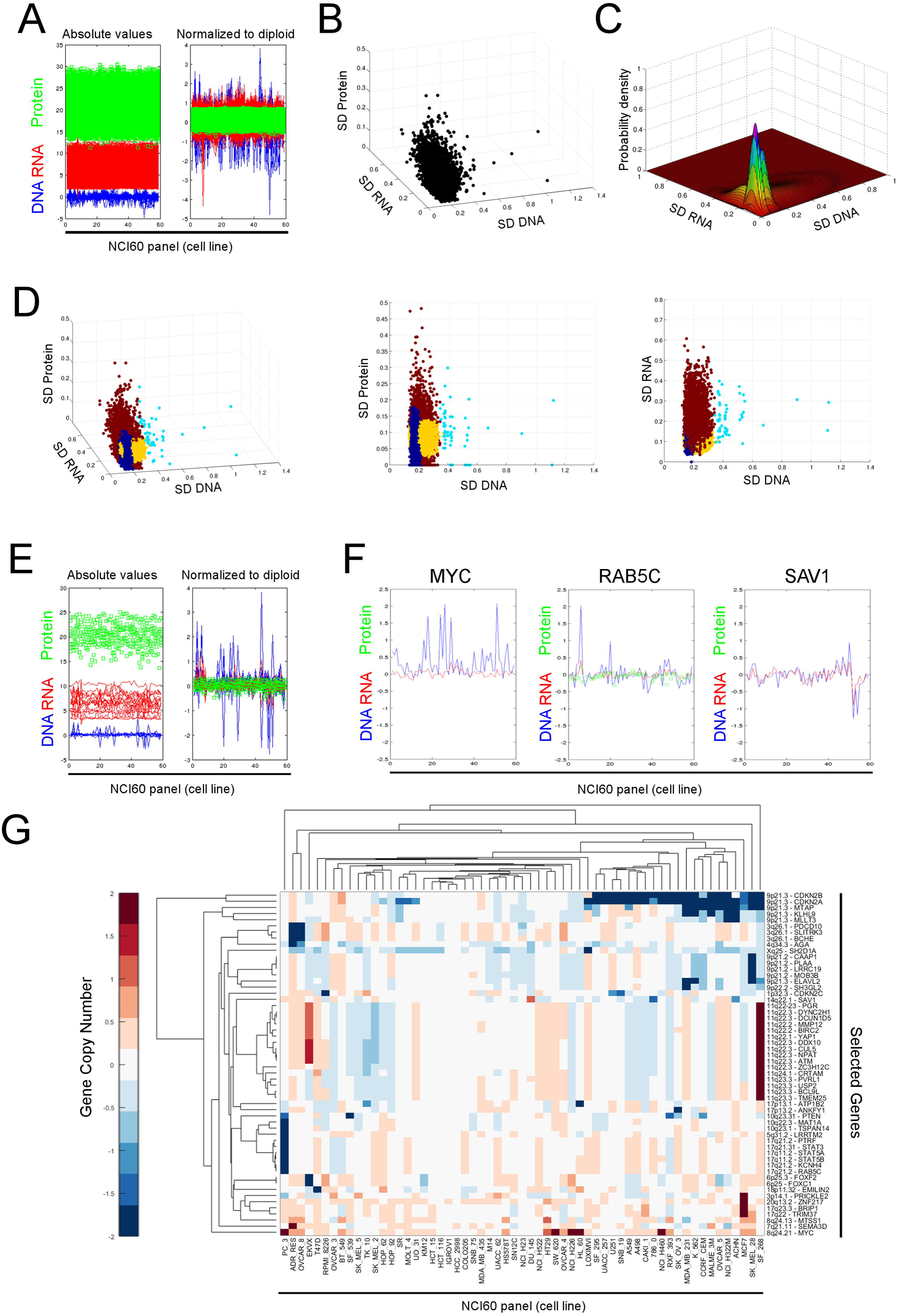
Identification of candidate genes under dosage compensation in the NCI60 panel. **A.** Input data of gene copy number (DNA, blue), gene expression (RNA, red) and protein levels (protein, green) of the NCI60 panel. The absolute values are shown on the left panel across the 59 cell lines and the right panel corresponds to the log2 values normalized to the averaged RNA and protein of the diploid cell lines for the respective gene (Normalized to diploid). **B.** Standard deviations (SD) of the DNA, RNA and Protein levels for each gene across the 59 cell lines of the NCI60 panel. **C.** Gaussian Mixture Model to identify a subpopulation of genes with high SD DNA and low SD RNA and/or low SD Protein (white arrow). **D.** Gene Clustering according to the model in C (left). The cyan cluster contains candidate genes under dosage compensation, characterized by high SD DNA, low SD Protein (middle) and low SD RNA (right). **E.** Absolute and Normalized values of selected candidate genes under dosage compensation. **F.** Examples of candidate genes under dosage compensation (*MYC* and *RAB5C*). **G.** Clusters of candidate genes according to their copy number variations across the 59 cell lines of the NCI60 panel.

To obtain a good separation of candidate compensated genes from the main cluster, we implemented a Gaussian mixture model (GMM) with these data (Figure 1C). After classification, we could observe one group of genes with high SD of DNA copies and relatively low SD of RNA and/or protein expression values (Figure 1D). This cluster showed the expected behavior with high amplitudes in DNA values but very low variation in RNA or protein expression (Figure 1D and Figure 1E). For instance, *MYC* presents high frequency of CNV amplifications in the NCI60 panel without the corresponding increase in RNA levels (Figure 1F left). A similar behavior is observed for *Rab5C* for both RNA and protein levels despite its copy number variation (Figure 1F middle), compared to a gene such as *Sav1* where copy number and RNA expression are well correlated (Figure 1F right). Next, we discarded those genes with orthologues in X/Y chromosomes since they cannot be differentiated using microarray techniques, and obtained a list of 56 gene candidates with putative dosage compensation at the transcript level. In some cases, there was also protein data confirming this behavior but there was no candidate with putative compensation at the protein level only. Moreover, we reduced our list of candidates by discarding genes with high DNA variation mostly due to deletions (more than 6 cell lines with compensated deletions) and kept only genes with at least 6 cell lines with compensated amplifications. For example, *MYC* presents 27 compensated amplifications within the NCI60 panel with very similar expression levels. This approach reduced our list to 21 candidate genes with dosage amplifications (Figure S1) potentially compensated at the transcript level across the NCI60 panel.

We next aimed to correlate the behavior of the copy number variations among these candidate genes to their corresponding chromosomal locations across the NCI60 panel. The clustergram in Figure 1G shows different clusters suggesting a similar behavior of copy number variations for these candidate genes. As expected, many of these genes are correlated by their genomic locations, especially gene clusters belonging to the chromosomal bands 3q26, 6p25, 8q24, 9p21-22, 10q22-23, 11q22-24, 17q11 and 17q21-23. These data suggest the existence of genes with low tolerance to variation in their RNA expression despite high copy number variation across the NCI60 panel, partially associated by common chromosomal locations. These genes are compensated candidates pointing to a transcriptional dosage compensation that propagates to the protein level. However, the protein data set of the NCI60 is limited, thus precluding the identification of candidates compensated at the protein level only.

### A mathematical model of gene dosage compensation mediated by a network of miRNA-transcription factor interactions for the NCI60 panel

We hypothesized, at this point, that the inhibition of a dosage compensation mechanism would release the brake imposed on the expression of all these additional copies leading to significant overexpression of these genes. If the dosage compensation of these genes was favored during cancer evolution to restrain the expression of key lethal genes, its overexpression could potentially lead to death of cell lines with these specific amplifications. Therefore, we focused our next steps on the description of possible mechanisms involved in dosage compensation of amplified genes identified in the NCI60 panel.

We asked whether dosage compensation mechanism could be mediated by systems-level properties arising from a complex regulatory network of gene expression. Since expression data of miRNAs (Blower et al., 2007) and transcription factors (TF) (Gmeiner et al., 2010) is available for the NCI60 panel, we performed a correlation analysis of the copy numbers of target genes with the expression levels of miRNAs and transcription factors in order to identify possible regulators responding to the dosage of our candidate genes (Figure S2). There are both miRNAs and TF with positive or negative correlation to the candidate copy numbers (Figure S2A), although most of those correlations do not correspond to the direct reported interactions (Figure S2B and Supplementary Section).

In order to model these interactions, we developed BioNetUCR, a computational platform to construct large scale models of miRNAs and TF interactions with the reverse approach, starting with a list of genes of interest that could be determined by differential expression analysis or customized by the researchers (Figure S3 (steps 1); see STAR methods, Network Construction (step 1)). Using this platform, we constructed a network of putative and validated regulatory interactions connecting all candidate genes with all their reported interactions with miRNAs and TFs and visualized it using CIRCOS table viewer (Krzywinski et al., 2009) (Figure S2C). Moreover, from the construction of this network topology, it is easy to infer that this network could sense changes in copy number of target genes only if they also have a TF-function (red colored interactions, Figure S2C). Only a gene candidate with TF function will trigger a signal that can be propagated throughout the network and back if this gene has a regulatory loop. This loop would compensate gene expression in response to changes in gene dosage.

Therefore, we reduced the network complexity to include only the NCI60 target genes with TF-function as they are the only genes for which a copy number variation can be sensed (*MYC*, *ZNF217, STAT3, FOXC1, FOXF2, PGR, STAT5A, STAT5B*). This resulted in a network of 517 nodes and 44016 arcs of putative and validated interactions. Since we collected about 450,000 experimentally validated interactions out of HTRI, Pazar, Transmir and Mirtarbase, we decided to include only experimentally validated interactions to further reduce model complexity. This led to a network of 78 nodes and 578 arcs of experimentally validated interactions. After this simplification, several target genes became dead ends, since no validated interactions are reported influencing other nodes of the network. Therefore, we removed them (BioNetUCR: DELETE SINK NODES, DELETE SOURCE NODES and DELETE AUTOREGULATED NODES), further reducing the model to include only 4 target genes with TF-function and validated interactions influencing other nodes of the network: *MYC*, *STAT3*, *STAT5A* and *STAT5B*. This led to a network of 65 nodes and 506 arcs. The nodes include 45 TFs and 20 miRNAs. In order to examine the target/miRNA/TF network for the presence of regulatory motifs with systems-level properties, we searched for positive and negative feedback loops (between miRNAs and TFs), coherent feedforward loops and incoherent feedforward loops. For this network, the arcs formed 16 feedback loops, 28 coherent feedforward loops and 45 incoherent feedforward loops (Figure 2A).

**Figure 2.**
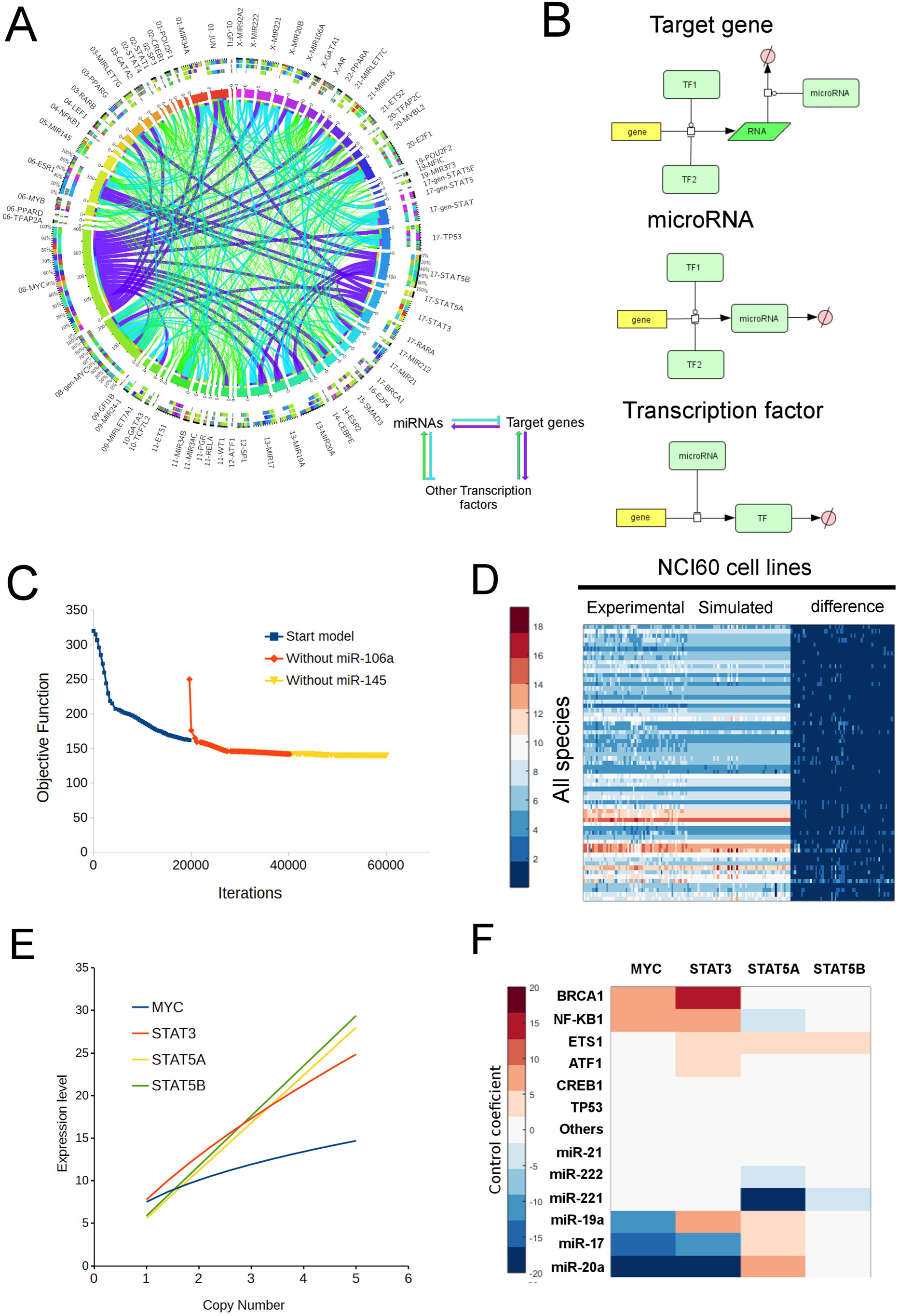
A mathematical model driven by NCI60 data leads to a phenotype of gene dosage compensation for *MYC* and *STAT3*. **A.** Network of miRNA-TF interactions for *MYC*, *STAT3*, *STAT5A* and *STAT5B*. **B.** Schematic representations of the modeled interactions for the target gene, miRNAs and associated TFs for ODE model construction. State transition arrows indicate synthesis or degradation, catalysis lines (circular-head) indicate activation and flat-headlines indicate inhibition. **C.** Results of the parameter estimation task to fit the ODE mathematical model to the experimental data upon systematic removal of the species with worse fitting and lowest control on the candidate genes, leading to a reduction in the objective function (difference between modeled and experimental data). **D.** Graphical comparison of modeled and experimental data shows no significant differences for the final resulting model. **E.** Results of parameter scan on the values of copy number for the candidate target genes showing gene dosage compensation for *MYC* and *STAT3*. **F.** The sensitivity analysis points to some candidate regulatory molecules controlling the concentrations of *MYC* and *STAT3*.

Thus, the regulatory motifs with potential systems-level properties to mediate dosage compensation are widely present within this putative network. However, the network topology alone is not enough to identify robust targets controlling the phenotype of interest since a gene-dosage response mediated by one or several of these regulatory loops will depend on the strength of the interactions (i.e. their kinetic parameter values). Due to the high complexity of this miRNA/TF regulatory network, we envisaged the construction of a large-scale mathematical model in order to gain insight into a possible mechanism of gene dosage compensation mediated by any of the many regulatory loops identified in this network. Therefore, beyond identifying a network, we used our platform to establish a full dynamical system based on ordinary differential equations (ODEs) (Figure S3A, step 3), which can be calibrated with existing experimental data (Figure S3A, step 4). To do so, we introduced ODE expressions for the schematic representations of the modeled interactions for the target gene, miRNAs and associated TFs (Figure 2B, Figure S3A step3) and translated them into differential equations. Reactions were represented using simple kinetic functions. Most follow mass action kinetics, except for transcription of genes that are induced or repressed by transcription factors, where linear activation or inhibition terms are added, as well as the degradation of RNA species that are affected by miRNAs, where linear inhibition terms were included. Gene copy numbers are also included as part of the mass action rate constant (See STAR Methods: Conversion of network to ODE model (Step 3)). The resulting biochemical model includes 65 species, 130 reactions and 305 parameters.

A quantitative systems-level approach for the calculation of the goodness of fit of a mathematical model describing the interactions enabled us to assess the feasibility of a proposed network topology for gene dosage compensation. The 305 model parameters included 65 copy numbers that were set as independent values and 240 kinetic parameters for fitting (degrees of freedom) that were estimated to fit the model to the 3,835 experimental data points (dependent variables) for a ratio of 16X. The model fitting was performed using the Parameter Estimation function of COPASI with the Hooke-Jeeves algorithm of optimization (Hooke and Jeeves, 1961) under the assumption that each cell line of the NCI60 panel represents a different steady state of this model depending only on the copy numbers of the genes in the network (STAR methods: BioNetUCR (step 5): Model parameter estimation using COPASI). We increased the weight of the objective function of the expression of the 4 target genes in order to favor the search of model parameter values leading to dosage compensation. Model refinement was performed by interrupting the fitting process after several days without any significant decrease in the objective function. At these periods of time, we calculated the goodness-of-fit for each species and performed sensitivity analysis using the sensitivity analysis framework of COPASI. Finally, we evaluated the model for dosage compensation by performing a parameter scan on the copy number (from 1 to 5) of the candidate genes under dosage compensation (STAR methods: Assessment of gene dosage compensation behavior and other simulations).

Using this network, we constructed and fitted a first ODE mathematical model, which was not able to reproduce any behavior of dosage compensation. Despite the initial set-backs, the turning point toward the modeling of the gene dosage compensation was the insight of the sensor loop: the group of interactions starting in a transcription factor and returning after a defined number of interactions with other species including miRNAs and other TFs. These include feedback and feedforward loops but also more complex interactions. The sensor loop itself also helps the network to adapt to fluctuations of the genes by repressing its expression. Hereby, we enriched our network for only those arcs establishing feed-back or feed-forward loops obtaining smaller models suitable for a faster parameter estimation (STAR methods: BioNetUCR (step 2): Optional Network Simplifications and Sensor Loops).

After constructing the new ODE model enriched in Sensor Loops, we performed a new parameter estimation and proceeded with a systematic elimination of nodes with the worse fitting and without control over the genes of interest (STAR methods: BioNetUCR (step 6 and step 7): Sensitivity analysis and removal of non-fitted/weak regulators). First, we calculated the p-values (paired t-Student to compare simulated with experimental data) for the goodness-of-fit for the resulting model, which presents a good fit for most species except for miR106a (p = 0.012) and no control of miR106a on any of the candidate genes. After removing miR106a, the resulting model achieved a lower objective function value (better fit) but miR145 had the worse goodness-of-fit (p=0.075) without significant control on the target genes. We continued by removing miR145 from the model to construct a new network and continued the parameter estimation to obtain an optimized model with a lower objective function (Figure 2C). For this last fit, there was no statistical difference of the values of the simulated species compared to the experimental data (p>0.1, Figure 2D), obtaining thereby a mathematical model describing the experimental data of our target candidate and their associated miRNAs and TFs.

Furthermore, we evaluated this model for gene dosage compensation by performing a parameter scan using COPASI for the copy number of the 4 target genes individually (STAR methods: Assessment of gene dosage compensation behavior and other simulations). As shown in Figure 2E, increasing the copy number of *MYC* from 1 to 5 leads only to an increase of 1.95-fold in *MYC* expression. For *STAT3*, an increase from 1 to 5 in copy number leads to an increase of 3.19-fold in *STAT3* expression. This was not the case for *STAT5A* and *STAT5B*, which copy number increases were not compensated in the model. We next performed a metabolic control analysis using COPASI to identify candidate miRNAs or TFs with high positive or negative regulatory control on the expression level of the 4 target genes. We plotted the 6 miRNAs with the highest negative control and the 6 TFs with the highest positive control (Figure 2F). The results showed that *MYC* levels are mainly controlled by an interplay between miR17, miR20a, miR19a and *BRCA1* and *NF-KB1*. *STAT3* levels are mainly controlled by miR17, miR20a, miR19a and *BRCA1*, *NF-KB1*, *Ets1* and *ATF1*. Although *STAT5A* and *STAT5B* were not compensated upon changes in their copy numbers, they also present control in their expression levels by some of the same miRNAs and TFs, suggesting that the mechanism of *MYC* and *STAT3* compensation could also impact the levels of *STAT5A* and *STAT5B*. These observations demonstrate that our mathematical model is able to describe a theoretical network of validated interactions between miRNAs and TFs with systems-level properties mediating direct dosage compensation of *MYC* and *STAT3*, with a possible impact on *STAT5A* and *STAT5B*. If this model is correct, the inhibition of some miRNAs negatively controlling the expression levels of the target genes could potentially block gene dosage compensation.

### A minimal model of MYC and STAT3 dosage compensation leads to the identification of a putative mechanism of gene-dosage compensation in aneuploid cancer

We next asked which are the minimal components of the network to identify the exact regulatory loops mediating gene dosage compensation and to design strategies to interfere with this mechanism *in silico*. First, we sought to identify the essential species involved in the feedback or feedforward loops controlling gene dosage compensation. We performed parameter scans changing the copy number of the compensated genes (*MYC* and *STAT3*) and observed the behavior of the concentrations of the miRNAs and TFs of the model. For *MYC* copy number scan, the *MYC* expression showed a compensated increase, observed also for several TFs and miRNAs (Figure 3A left), including those identified by metabolic control analysis (miR-19a, miR-20a, miR-17, Figure 2F) but others, such as *STAT3* and miR21, decreased with increasing *MYC* copy numbers (Figure 3A left). For *STAT3*, increasing gene copy numbers led to a compensated increase of *STAT3*, several miRNAs including miR21, miR17, miR221, miR19a and miR20a and other TFs but also a decrease in *STAT5A*, *ETS1* and *PPARG* (Figure 3A right).

**Figure 3.**
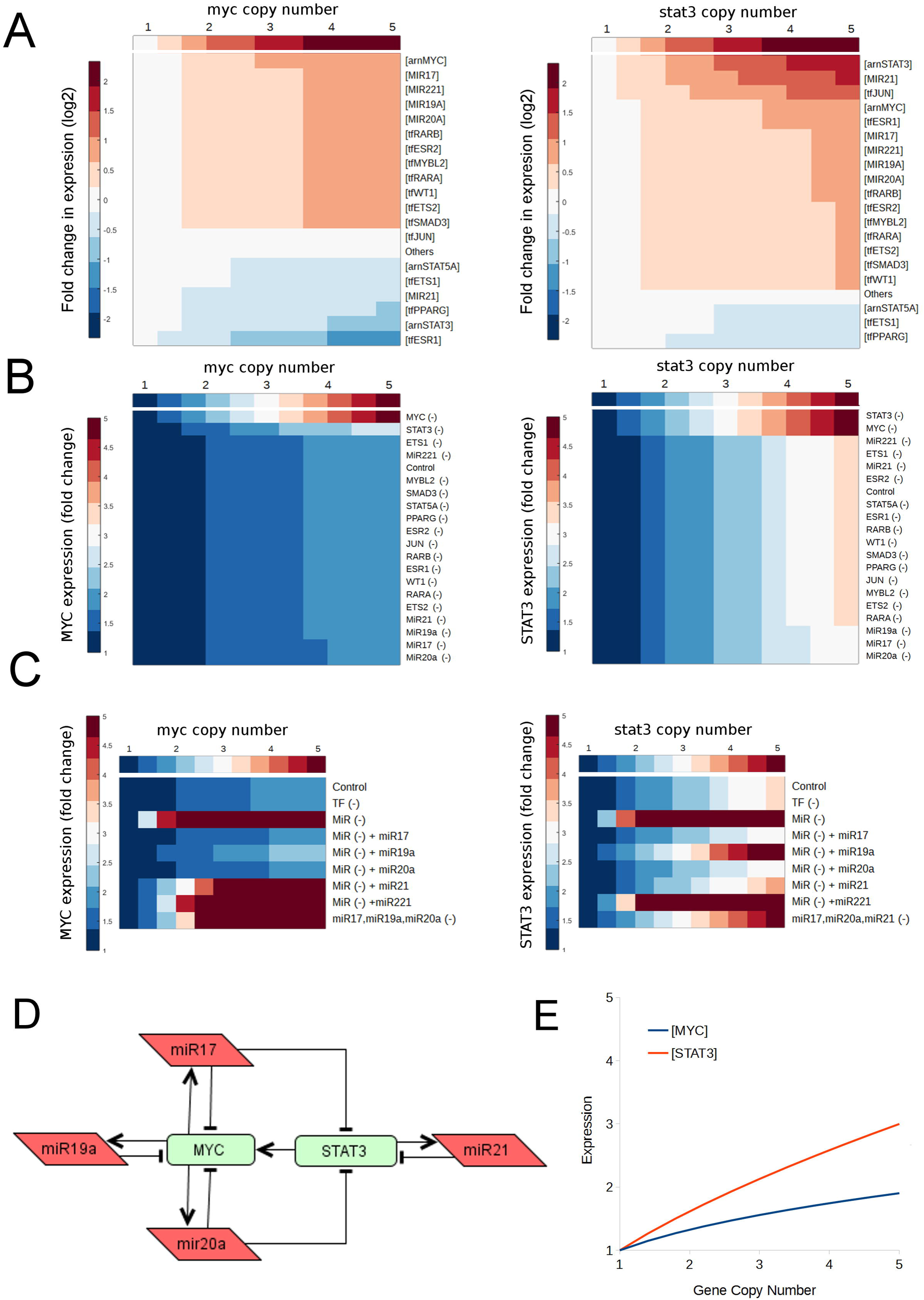
Biology-inspired experiments with the mathematical model lead to the identification of a minimal model of gene dosage compensation for *MYC* and *STAT3*. **A.** A parameter scan on *MYC* (left) and *STAT3* (right) copy numbers identifies which molecules increase (red) or decrease (blue) together with varying copy numbers of those two genes. **B.** The single inhibition of those varying molecules has no effect on the parameter scan to evaluate gene dosage compensation of *MYC* (left) and *STAT3* (right). The expression remains blue similar to control conditions except for the inhibition of *MYC* or *STAT3* themselves. **C.** The inhibition of groups of molecules and the restoration of single miRNAs indicate a redundant mechanism for the gene dosage compensation of *MYC* (left) and *STAT3* (right). **D.** A minimal model of gene dosage compensation for *MYC* and *STAT3* is hypothesized based on the results of these experiments (simplified representation). **E.** The fitting and parameter scan demonstrates that this minimal model recapitulates the dosage compensation of *MYC* and *STAT3*.

To assess the role in dosage compensation of these miRNAs and TFs responding to the changes in *MYC* and *STAT3* copy numbers, we performed single *in silico* inhibitions of the corresponding nodes by setting their interaction (inhibition/activation) parameters to 0. The inhibitions of single species altered the steady state levels of expression of *MYC* and *STAT3* for the basal conditions (copy number of 1) but this effect is not shown due to the normalization of the expression, to better appreciate the effect on gene dosage compensation (Figure 3B). The parameter scans under these conditions showed that *MYC* and *STAT3* compensations are completely abolished when their respective interaction parameters are set to 0, confirming that their TF function is essential for the network to sense the changes in copy number (*MYC*(-) and *STAT3*(-) conditions, Figure 3B). Also, *MYC* compensation is slightly altered when *STAT3* is inhibited (*STAT3*(-)) and *STAT3* compensation is completely abolished when *MYC* is inhibited (*MYC*(-)), suggesting an interplay in the mechanisms of gene dosage compensation of these two genes. In contrast, the single inhibition for all remaining TFs and miRNAs had little or no effect on *MYC* and *STAT3* dosage compensation, suggesting that this mechanism could be redundantly mediated by more than one miRNA or TF.

Therefore, we performed another *in silico* experiment changing the copy number of *MYC* and *STAT3* while all miRNAs or TFs were inhibited, except by *MYC* and *STAT3* themselves (Figure 3C). The results showed that the inhibition of all the other TFs (TF (-)) had no effect on the dosage compensation of *MYC* and *STAT3*. However, the inhibition of all miRNAs (MiR(-)) completely abolished the dosage compensation for both genes and led to a huge increase in their expression. We then proceeded to restore some miRNAs individually back into the MiR inhibited model, for example (MiR(-) + miR17) to simulate the effect of the active miR17 alone. Surprisingly, more than one miRNA was able to restore dosage compensation for both *MYC* and *STAT3*, confirming thereby that this mechanism is redundant. Indeed, miR-17, miR-19a and miR20a were individually able to compensate *MYC* dosage and miR17, miR20a and miR21 were able restore *STAT3* dosage compensation (Figure 3C). Additionally, to confirm the role of these redundant miRNAs on gene dosage compensation *in silico*, we performed their triple inhibition in the complete model. Indeed, the triple inhibition (miR17, miR19a and miR20a (-)) was able to block *MYC* dosage compensation (Figure 3C left), whereas (miR17, miR20a, miR21 (-)) abolished *STAT3* dosage compensation (Figure 3C right), confirming that *MYC* and *STAT3* dosage compensation is mediated by a redundant mechanism.

Based on these results of redundant compensation and the interplay between *MYC* and *STAT3*, we proposed a minimal network topology of gene dosage compensation for *MYC* and *STAT3* (Figure 3D). The Systems Biology Graphical Notation (SBGN) representation of this model is presented in Figure S3B. After reconstructing the interactions from the original network, the minimal network states that *MYC* is compensated by 3 redundant negative feedback loops formed with miR17, miR19a and miR20a. The compensation of *STAT3* is mediated by 1 feedback loop with miR21 and 2 feedforward loops: (*STAT3*-*MYC*-miR17-*STAT3*) and (*STAT3*-*MYC*-miR20a-*STAT3*). From this minimal network topology, we constructed a minimal mathematical model of gene dosage compensation for *MYC* and *STAT3*, including only 18 parameters against 354 experimental data points (20× ratio). After fitting this minimal model to the experimental data and repeating the parameter scan experiments for gene copy numbers of *MYC* and *STAT3*, we reconstituted the same dosage compensation mechanism observed in the original large model of interactions (Figure 3E).

These results indicate that we identified *in silico* a minimal model of gene dosage compensation for *MYC* and *STAT3* mediated by their interactions with miR17, miR19a, miR20a and miR21. Indeed, the sensitivity analysis and the *in silico* experiments enabled us to identify the most robust target nodes to regulate the phenotype of interest, to determine the mechanism behind that phenotype and the interventions on those target nodes. This results in a smaller model with a higher explanatory power than the original (large) network.

### MYC is widely compensated across different types of cancer

To assess the extent of *MYC* and *STAT3* dosage compensation across different cancer types in a larger dataset, we used the same criteria of the NCI60 study to identify candidate genes under dosage compensation for 898 cell lines of Cancer Cell Line Encyclopedia (CCLE) having both Expression and CNV data (STAR Methods: Model of *MYC/STAT3* dosage compensation with CCLE). First, the expression data was normalized to the mean of the diploid cell lines for each gene, and both the normalized expression and the absolute copy number was log2 transformed and centered to 0 (diploidy) for a direct comparison (Figure 4A, left). Likewise, the standard deviations (SD) of both log2 transformed data were calculated and plotted to search for clusters of putative candidate genes under dosage compensation. The scatter plot shows a cluster (blue) of 152 genes with the highest SD of CNV and relatively low SD in expression, indicating the presence of some genes with low tolerance to variation in expression despite of having the highest variation in their CNVs (Figure 4B). When the expression and CNV data of this cluster is plotted, a narrower amplitude of expression can be observed compared to the full data set as expected (Figure 4A, right). We could also locate our two main candidates of the NCI60 study within the CCLE data. *STAT3* has lower copy number variation compared to the NCI60 data set but a similar low variation in expression. In contrast, *MYC* could be mapped as one of the genes with the highest variation in CNV and the lowest corresponding variation (Figure 4B). A closer look at the individual *MYC/STAT3* data across CCLE shows that *MYC* has many more amplifications with high copy numbers, all of them probably compensated. *STAT3* in contrast, has few compensated amplifications of only one additional copy (Figure 4C). There are also several genes with higher variation in CNV compared to *MYC* and among them there is one with a particularly low variation in expression, corresponding to TBC1D3B (https://www.genecards.org/cgi-bin/carddisp.pl?gene=TBC1D3B). TBC1D3B codes for a protein involved in *RAB* GTPase signaling and vesicle trafficking, and located in chromosome 7q, a shared chromosomal location with some of our previous NCI60 candidates including *STAT3*, *STAT5A* and *STAT5B*. These findings confirmed that *MYC* is one of the most frequently amplified and compensated genes across CCLE according to our criterion to identify dosage-compensated candidates and there are other candidates under robust dosage compensation.

**Figure 4.**
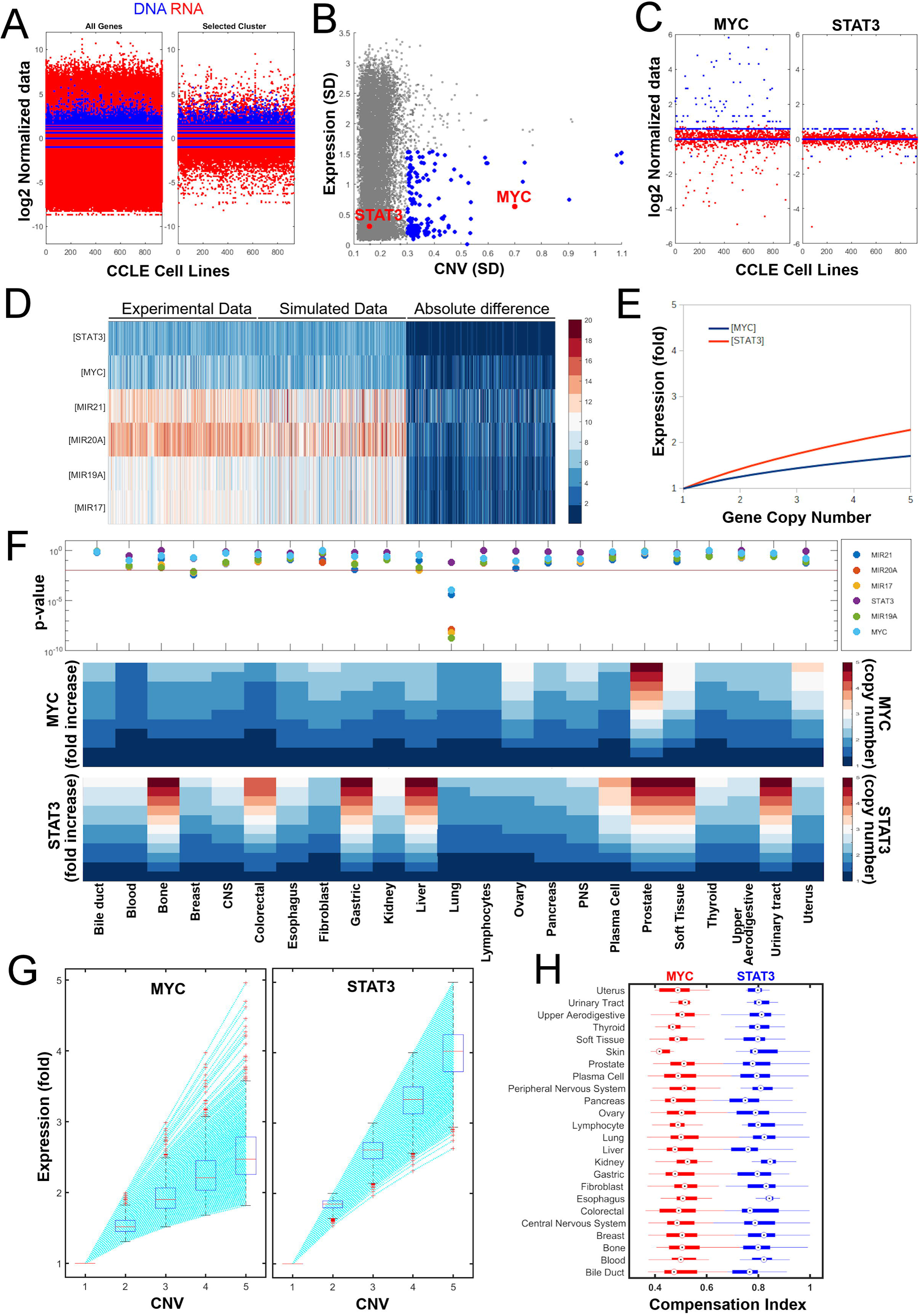
Status of *MYC* and *STAT3* compensation across the CCLE cell lines. **A.** Input data of gene copy number (DNA, blue) and gene expression (RNA, red) of CCLE normalized to diploid average (expression) and log2 transformed for all the genes (right) and the selected cluster in B (right). **B.** Standard deviations (SD) of the Copy Number Variation (CNV) and Expression for each gene across the cell lines of the NCI60 panel. A selected cluster of genes with low tolerance to changes in expression despite of high variability in Copy Number was detected (blue). The respective positions of *MYC* and *STAT3* are shown in red. **C.** Gene copy number (DNA, blue) and gene expression (RNA, red) of CCLE normalized to diploid average (expression) and log2 transformed for *MYC* and *STAT3.* **D.** Fitting results of Minimal Model of MYC/STAT3 dosage compensation compared to the experimental data set (p-values for miR21: 0, miR20a: 8×10^−10^, miR17: 1.6×10^−9^, *STAT3*: 0.126, *MYC*: 7×10^−8^). **E.** Parameter Scan for *MYC* and *STAT3* copy numbers of the pan-CCLE model shows gene dosage compensation for both *MYC* and *STAT3*. **F.** Results of parameter estimation for each of the species of tumor-type specific minimal models of dosage compensation (upper panel, p > 0.01). The corresponding parameter scans on gene copy numbers of *MYC* (middle panel) and *STAT3*(lower panel) display heterogeneity in gene dosage compensation. **G.** Single-experiment parameter scans of copy numbers (from 1 to 5) show the heterogeneity in *MYC* and *STAT3* dosage compensation. On each box, the central mark indicates the median, and the bottom and top edges of the box indicate the 25th and 75th percentiles, respectively. The whiskers extend to the most extreme data points not considered outliers, and the outliers are plotted individually using the ‘+’ symbol. **H.** Distribution of *MYC* and *STAT3* dosage compensation indexes by tumor type.

We set out to investigate whether this behavior of apparent dosage-compensation of *MYC/STAT3* could be explained by the same circuits postulated for the NCI60 data. We studied those circuits at 3 different levels. First, we tried to fit a pan-CCLE mathematical model using all the cell lines to fit the minimal model of *MYC/STAT3* dosage compensation. Although we could not fit the model for the different species (except for *STAT3*) a tendency can be observed when the simulated data set is compared to the experimental data (Figure 4D) and the resulting model recapitulates the behavior of gene dosage compensation for both genes (Figure 4E). Second, we hypothesized that the high heterogeneity of CCLE impairs a correct fitting of the minimal model and therefore decided to fit the minimal model to tumor-type specific datasets. We achieved a complete fitting for all the species of most tumor-specific models of gene dosage compensation except for Lung and Breast cancer (Figure 4F, upper panel). We next tested the tumor-type models for their capability to compensate by parameter scans on *MYC/STAT3* copy numbers and could observed a generalized *MYC* dosage compensation across tumor types with certain degree of heterogeneity, except for the prostate cancer model that showed no dosage compensation (Figure 4F, middle panel). In contrast, there was a much higher heterogeneity in *STAT3* dosage compensation across tumor types. Eight tumor-type specific models were not able to compensate *STAT3* dosage at all (Figure 4F, lower panel).

In order to further characterize this heterogeneity in gene-dosage compensation, we used the model as a tool to perform individual parameter estimation of single cell lines (see STAR methods: material and methods: Calculation of Compensation Index). With the single-experiment models, we performed parameter scans to calculate the increase in gene expression *in silico* at steady-state upon increasing gene copy numbers from 1 to 5. The values were normalized to the expression at CNV=1 for comparison. We could therefore observe that the majority of cell line-specific models has the capability of *MYC* dosage compensation in their current configurations (determined by their parameter values) with only few outliers, as shown by the sub-linear increases in gene expression upon an increasing CNV (e.g. an increase in a copy number from 1 to 5 leads to an increase of only 2.3 fold in expression; Figure 4G, left panel). In contrast, *STAT3* dosage compensation is much weaker for the majority of cell-line specific models with an average increase of 4-fold upon a CNV increase from 1 to 5 (Figure 4G, right panel). We therefore defined a compensation index for comparison, defined as the ratio of the normalized expression fold with a copy number of 5 divided again by 5. Thereby, a compensation index close to 1 indicates very weak compensation and a compensation index close to 0 indicates a strong compensation *in silico*. When we plotted the compensation indexes of the cell line-specific models across tumor types, we could observe that their parameter configurations enable a robust and wide compensation of *MYC* across tumor types, whereas *STAT3* is weakly compensated (Figure 4H). These results indicate that *MYC* dosage compensation is enabled in most tumor types despite the high level of heterogeneity observed in cancer. Although this heterogeneity and the high complexity of the CCLE dataset precludes the fitting of larger mathematical models of gene dosage compensation, it represents a very important tool to validate minimal models of gene regulation and quantify the degree of heterogeneity in emerging systems-level properties such as dosage compensation.

### The kinetic parameters of the TF-miRNA interactions determine the capability of a putative network motif for gene dosage compensation

To this point we were able to identify *in silico* a minimal model of gene-dosage compensation for *MYC* and *STAT3* mediated by their interactions with 4 miRNAs. However, there is heterogeneity in dosage compensation across tumor types and different degrees of compensation (e.g. the difference between the compensation index of *MYC* and *STAT3*). There, we next asked whether we could identify the quantitative properties of these network motifs conferring gene dosage-compensation. First, we designed a rationale for the identification of motifs mediating gene dosage-compensation based on our findings with *MYC* and *STAT3* (Figure 5A). To evaluate this rationale, we constructed individual networks of our remaining NCI60 candidates using the experimentally validated interactions included in BioNetUCR (*ZNF217, FOXC1, FOXF2, PGR*, *STAT5A*, *STAT5B*). Next, we looked for regulatory motifs of one, two and three nodes returning to the gene candidates. Only *PGR*, *STAT5A* and *STAT5B* had regulatory motifs returning to them after one, two or three nodes (not shown). Using these 3 networks, we built corresponding COPASI models and fitted them to the NCI60 dataset and evaluated the ability of the model to compensate by a parameter scan (Figure 5B). Afterwards, we performed *in silico* experiments by inhibiting individual or groups of model species (such as those experiments performed to identify the minimal model of *MYC*-*STAT3*) in order to identify minimal models for *STAT5A* and *STAT5B* dosage compensation. We identified an extended feedback loop for the dosage compensation of *STAT5A* and 3 extended feedback loops for the dosage compensation of *STAT5B* (Figure 5C). Indeed, those minimal models recapitulated the behavior of dosage compensation as expected (Figure 5D).

**Figure 5.**
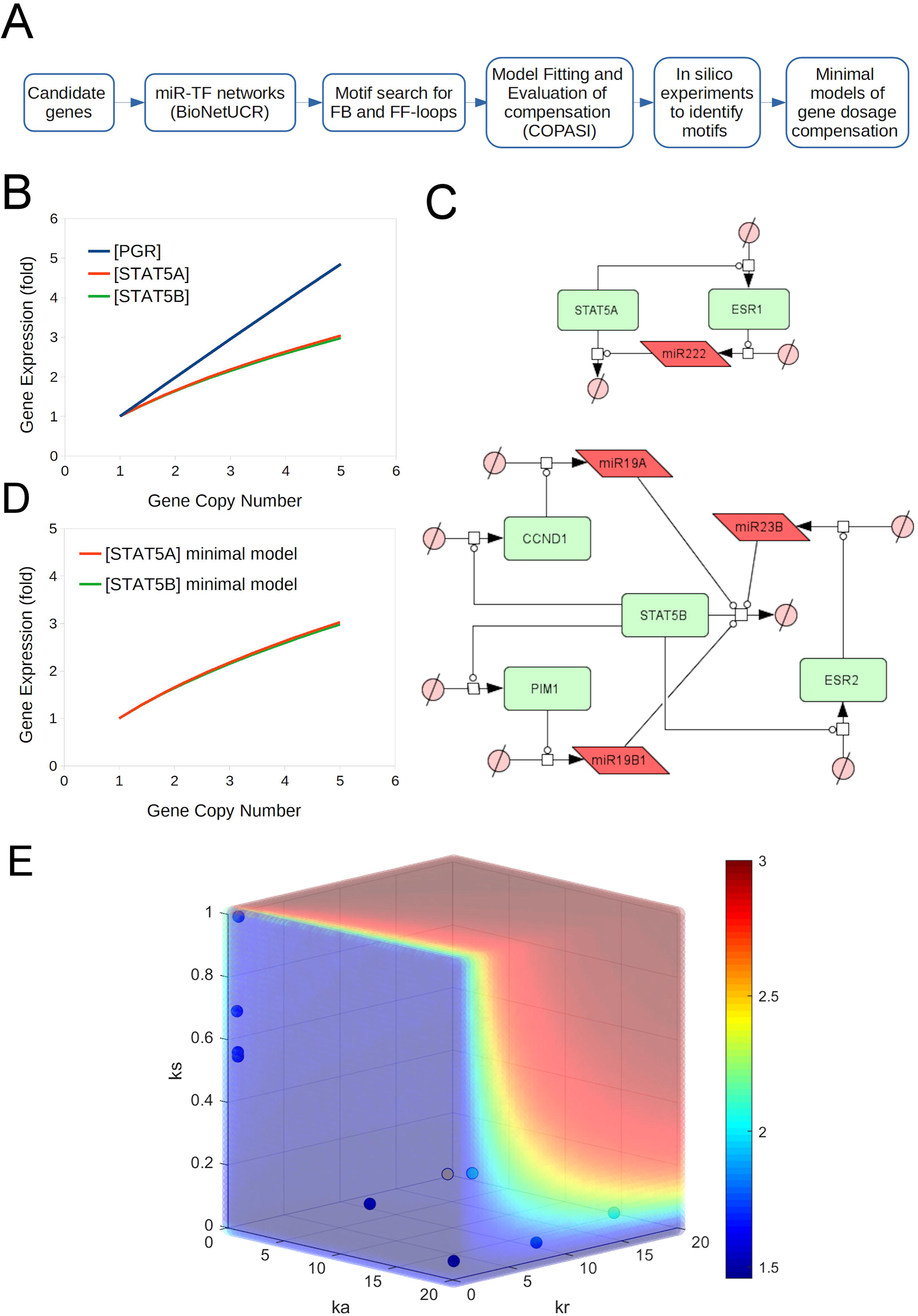
The gene dosage compensation depends on the kinetic parameters of the interactions of basic regulatory network motifs. **A.** Rationale of a biocomputational pipeline for the identification of motifs mediating gene dosage compensation. **B.** Evaluation of gene dosage compensation for the networks containing feedback and feedforward loops for PGR, *STAT5A* and *STAT5B*. **C.** The postulated network topologies of the minimal models of gene dosage compensation include 1 extended feedback loop for *STAT5A* and 3 extended feedback loops for *STAT5B*. State transition arrows indicate synthesis or degradation, catalysis lines (circular-head) indicate activation and flat-head-lines indicate inhibition. **D.** These minimal models recapitulate the gene dosage compensation for *STAT5A* and *STAT5B*. **E.** A triple-parameter scan on the values of ks (synthesis rate of the miRNA), ka (activation parameter of the TF) and kr (repression parameter of the miRNA) under the two different copy numbers of MYC (1 and 3). The ratio between these two scans is presented to display the three-dimensional landscape of gene dosage compensation and its dependence on the values of these kinetic parameters of some basic regulatory network motifs. Individual cases of regulatory motifs related to dosage compensation are located within the compensated region (blue) of this regulatory landscape.

To study the kinetic properties of the regulatory loops of *MYC*, *STAT3*, *STAT5A* and *STAT5B* postulated hereby as dosage compensation circuits, we explored the kinetic parameters describing the interactions between miRNAs and TFs by performing combined parameter scans in COPASI using the minimal model of *MYC*-*STAT3*. Since the compensation in that model is redundant, we inhibited miR19a and miR20a so that we can evaluate only the behavior of the single negative feedback loop between *MYC* and miR17. Next, we monitored the concentration of *MYC* for two conditions: a *MYC* copy number of 1 and a *MYC* copy number of 3. Under these two conditions we scanned the effect of different values of the parameters and found that 3 parameters were best to separate the parameter landscape into two regions, a non-compensation space and a compensation space of parameters: the ks (synthesis rate of the miRNA), ka (activation parameter of the TF) and kr (repression parameter of the miRNA). After calculating the ratio of *MYC* concentrations of those 2 conditions, we obtained a three-dimensional landscape of gene dosage compensation with a color-coded heatmap of the fold increase in *MYC* expression upon a 3-fold increase in copy number (Figure 5E). The blue region corresponds to the dosage-compensation space of parameters whereas the red region represents the space of parameters leading to a linear increase of gene expression as a function of copy number (no compensation). Moreover, when we located the positional values of the parameters corresponding to the compensating motifs, they all appeared in the blue region of compensation (Figure 5E) except by the interactions involving *STAT3* that appeared close to the green/yellow transition zone. These results confirmed that the simple identification of network motifs such as feedback or feedforward loops is not enough to identify systems-level properties such as gene-dosage compensation. This behavior is indeed dependent on the data-driven identification of the magnitude of the parameter values accounting for the emergence of this property of gene dosage compensation.

### A genetic tug-of-war approach enables the experimental demonstration of MYC dosage compensation

Although our proposed minimal model of gene dosage compensation was based on reported, experimentally-validated interactions and the interaction parameters were fitted to experimental data, it was necessary to confirm the ability of those regulatory circuits to perform gene dosage compensation. Therefore, we undertook the experimental validation of the *MYC* dosage compensation circuits by designing a novel tug-of-war approach to identify gene dosage compensation at the transcriptional level, inspired by the approach of Ishikawa et al for the identification of post-translational dosage compensation (Ishikawa et al., 2017). In this report, the authors inserted a TAP tag next to the endogenous target gene to be evaluated. When the protein was over-expressed by transfecting a plasmid with an exogenous version of the gene, they observed downregulation of the endogenous protein, thus demonstrating dosage compensation at the translational level.

We developed an approach to identify gene-dosage compensation at the transcriptional level based on the same principles: 1. To differentiate the exogenous and endogenous transcripts of *MYC*, we overexpressed an exogeneous version of *MYC* ((exo)*MYC*) with a differential codon optimization compared to the human endogenous *MYC* ((endo)*MYC*. 2. We removed the 3’UTR segment of the exogenous *MYC* to avoid any miRNA regulation on this transcript so that it can exert a higher pressure on the *MYC* regulatory circuits. 3. Although both transcripts will be translated into the same protein and exert the same regulatory activities, they can be differentiated at the transcript level by RT-qPCR using specific primers and probes. Under the pressure of the dosage compensation circuits, we hypothesize that the exogenous *MYC* overexpression will lead to the down-regulation of endogenous *MYC*.

First, we transfected NCH82 glioblastoma cells with either a control plasmid containing GFP (GFP-Ctrl) or a plasmid containing GFP and the exogenous *MYC* (GFP-(exo)*MYC*) (Figure 6A). We validated the ability of our sets of primers and probes to differentiate between the endogenous and exogenous *MYC* transcripts. The expected band of 404 bp for endogenous *MYC* (Endo-) was observed in untransfected cells (control) and in the GFP-Ctrl and (exo)*MYC*-GFP transfected cells when detected with the corresponding primers and probe designed for the endogenous *MYC* transcript. The band of 216 base pairs expected for the exogenous *MYC* (Exo-) was observed only in the (exo)*MYC* transfected cells when detected with the primers and probe designed for the exogenous *MYC* (Exo) (Figure S4A).

**Figure 6.**
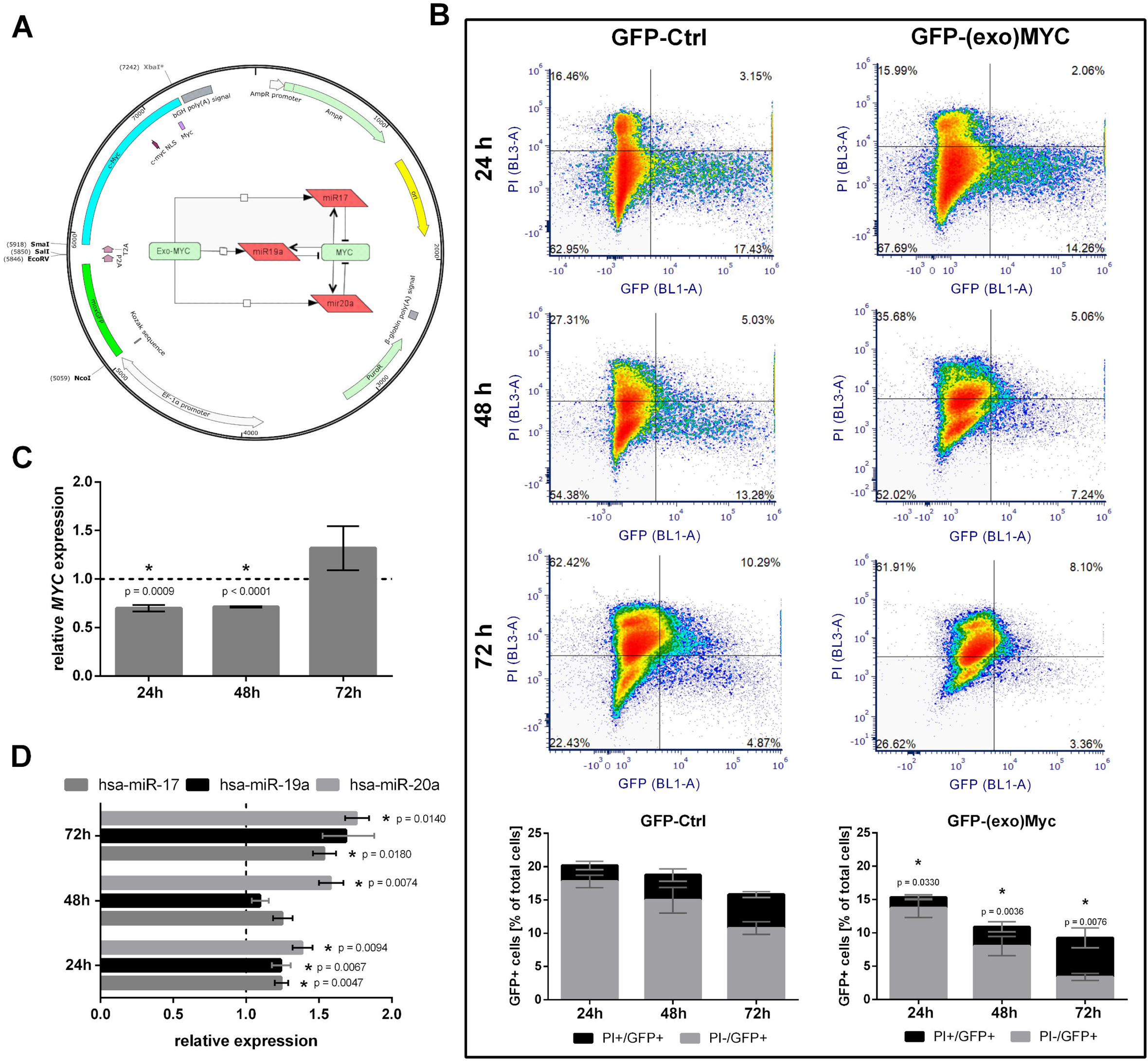
Overexpression of exogeneous *MYC* decreases endogenous *MYC* and increases expression of the predicted miRNA in human glioblastoma NCH82 cells. **A.** Vector map of the construct pSBbi-moxGFP-P2A-T2A-c*MYC*-Puro for the expression of (exo)*MYC* with a deletion of the 3’UTR region to avoid miRNA regulation to induce miRNA overexpression and the subsequent down-regulation of endogenous *MYC*. **B.** Upper part: Representative dot plots showing PI and GFP intensities of transfected NCH82 cells 24, 48 and 72 h after transfection. Left panel: cells transfected with the control GFP-plasmid (GFP-Ctrl); right panel: cells transfected with the (exo)*MYC* plasmid (GFP-(exo)*MYC*). Quadrants from upper left to lower left: Q1: PI+/GFP-; Q2: PI+/GFP+; Q3: PI-/GFP+; Q4: PI-/GFP-. Lower part: Statistical analysis of flow cytometric measurements of the PI+/GFP+ and PI-/GFP+ populations of transfected cells. Marked with * are statistically significant changes of total GFP+ cells in the GFP-(exo)*MYC* transfected cells compared to GFP-Ctrl transfected cells. (n = 6); **C.** Quantitative RT-PCR detection of endogenous *MYC* expression in the PI-/GFP+ population of GFP-(exo)*MYC* transfected cells relative to the PI-/GFP+ population of GFP-Ctrl transfected cells. (n = 3); **D.** Quantitative RT-PCR detection of hsa-miR-17, hsa-miR-19a and hsa-miR-20a in the PI-/GFP+ population of GFP-(exo)*MYC* transfected cells relative to the PI-/GFP+ population of GFP-Ctrl transfected cells. (n = 3); t-tests were performed to analyze means of samples with a significance defined by an α of 0.05. The resulting p-values were corrected for multiple comparisons using the Holm-Sidak method.

The efficiency of transient transfection is usually low, resulting in a subpopulation of transfected cells hindering the interpretation of any quantitative assay to monitor gene expression out of whole-population lysates. Therefore, we proceeded to monitor the GFP expression and cell death by flow cytometry (Figure 6B) and fluorescence microscopy (Figure S4B). We noticed that GFP-Ctrl transfected cells conserved a similar intensity in their fluorescence over time as shown by the fluorescence microscopy images. In contrast, (exo)*MYC*-GFP transfected cells started with a dim GFP fluorescence that did not increase over time. In fact, the number of transfected cells decreased over time and some of these cells showed a rounded morphology, suggesting that exogenous *MYC* overexpression is cytotoxic (Figure S4B). To quantify and confirm these results, we performed flow cytometry experiments to monitor GFP fluorescence and cell death, using propidium iodide (PI) staining. As shown in Figure 6B, the condition of GFP-Ctrl transfected cells displayed 20% GFP positive cells including 2% of dead cells at 24 h post-transfection. This amount decreased to 16% with 4% of dead cells at 72 h post transfection. The condition of (GFP-(exo)*MYC)* transfected cells showed a significant lower efficiency of transfection (15% of GFP positive cells after 24 h of transfection). This value decreased to 9% at 72 h post transfection with a high proportion of dead cells (more than 50% of the transfected cells), supporting our suggestion that *MYC* overexpression is cytotoxic.

In order to assess if *MYC* is dosage-compensated at the transcriptional level in the subpopulation of transfected NCH82 cells, we performed cell sorting of the viable (PI negative) GFP positive population of cells 24 h, 48 h and 72 h post-transfection. We extracted RNA from the sorted subpopulations to confirm expression of exogenous *MYC* by RT-qPCR (Figure S4C). We observed a strong increase of exogenous *MYC* 24 h and especially 48 h post-transfection, but no difference compared to the GFP negative subpopulation 72 h post-transfection. To quantify the effects of exogenous *MYC* on the expression of endogenous *MYC,* we compared the expression of endogenous *MYC* in the PI-/ GFP+ population of GFP-Ctrl and GFP-(exo)*MYC* transfected cells (Figure 6C). Overexpression of *MYC* led to a significant reduction of approximately 30% of endogenous *MYC* expression at 24 h and 48 h post-transfection. 72 h after transfection we could not detect a significant difference in (exo)*MYC* expression (Figure S4C), neither in endogenous *MYC* expression (Figure 6C), probably caused by an exo-*MYC* induced cytotoxicity, resulting in the time-dependent death of *MYC* overexpressing cells. These findings confirm the existence of regulatory circuits modulating the expression of *MYC* through an increase in gene-dosage, indicating that *MYC* dosage compensation is an active process in NCH82 cancer cells. To further study the nature of these regulatory circuits, we evaluated the expression of the 3 miRNAs predicted by the minimal model of *MYC* dosage compensation. As expected from the model, miR-17, miR-19a and miR-20a expression did increase upon (exo)*MYC* overexpression, suggesting that those miRNAs are involved in the regulatory circuits of *MYC* dosage compensation (Figure 6D).

### The blockade of MYC gene dosage compensation induces MYC dosage-dependent cytotoxicity

We next asked whether the blockade of *MYC* dosage compensation induces cell death and if the gene dosage plays a role in cell sensitivity to this blockade. Since the minimal model indicates that *MYC* dosage compensation is redundantly regulated by miR17, miR19a and miR20a, we first explored *in silico* the properties to identify the optimal strategies to achieve an inhibition of gene dosage compensation leading to *MYC* overexpression. For that purpose, we used the minimal model of *MYC* compensation to perform a triple parameter scan on the miRNA synthesis parameters (ks) to systematically reduce the amounts of the 3 miRNAs and identify steady states leading to *MYC* overexpression. In addition, we aimed to evaluate the effect of *MYC* dosage on *MYC* overexpression under the inhibition of gene dosage compensation. Therefore, the scan was repeated under 3 conditions of different *MYC* copy numbers (2, 4 and 7 corresponding the *MYC* copy numbers of 3 colon cancer cell lines of the NCI60 panel, see below) and the effect on the *MYC* concentration was observed. Although there are some slight alterations under a complete depletion of the 3 miRNAs for the diploid condition (CN = 2), the increase in *MYC* expression is much higher with increasing copy numbers, and less miRNA depletion is required to trigger the increase in *MYC* expression (Figure 7A). The results indicate that the interruption of *MYC* dosage compensation requires the blockade of the 3 miRNAs, although less inhibition of miR17 is required compared to miR19a and miR20a. Moreover, the *MYC* overexpression was easier to achieve with higher *MYC* copy numbers. These in silico results indicate that the perturbation of the mechanism of dosage compensation induces a gene dosage-dependent increase in *MYC* expression and suggests that *MYC*-amplified cells will be more sensitive to the blockade of *MYC* dosage compensation (Figure 7A).

**Figure 7.**
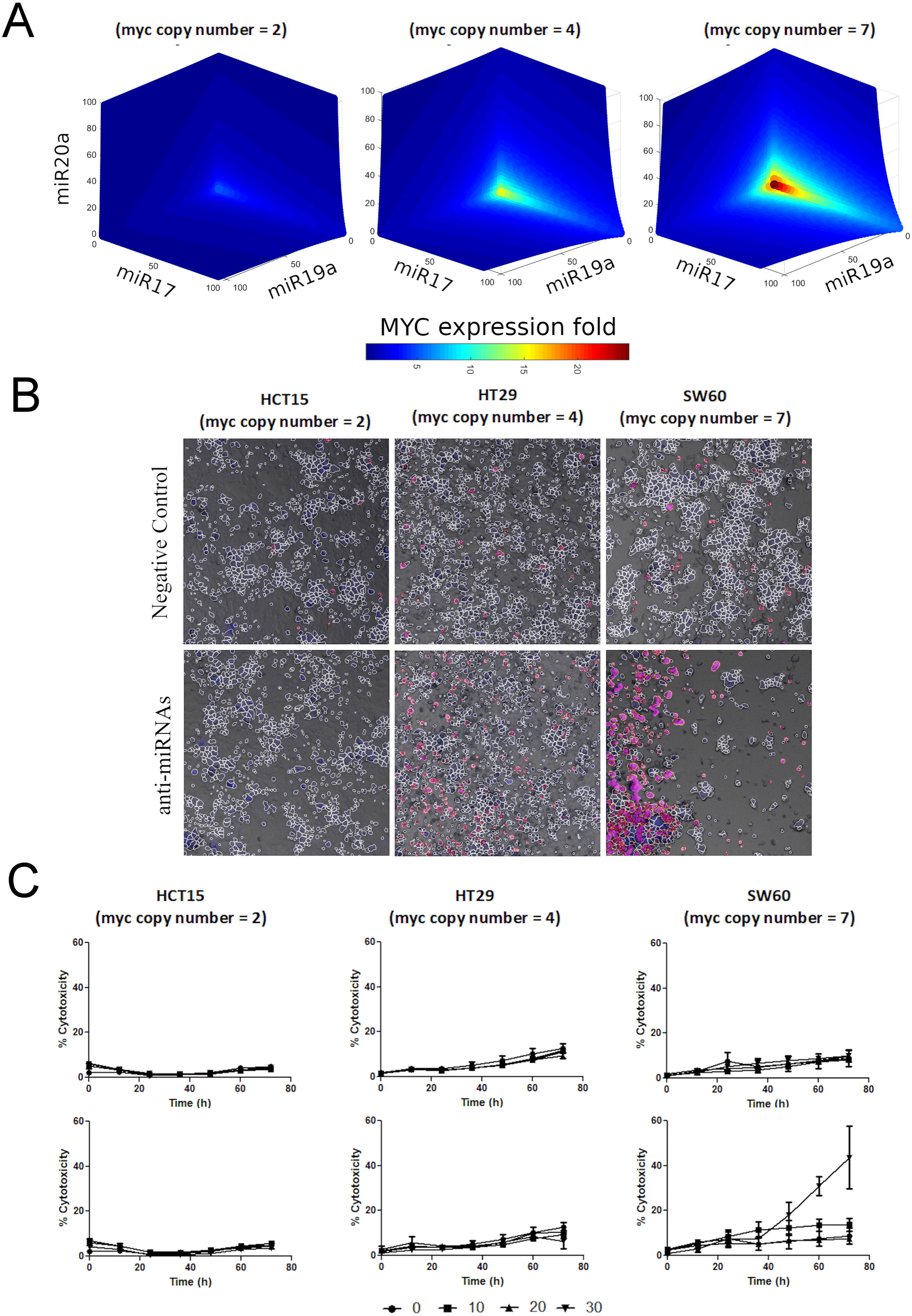
The inhibition of *MYC* dosage compensation induces dosage-dependent cytotoxicity. **A.** *In silico* simulations for the dependence of *MYC* concentration on the decreasing amounts of the 3 compensating miRNAs. The steady-state simulated MYC expression fold increase (compared to unperturbed model where MYC=1) in function of the percentage of the respective miRNAs (x,y,z) shows that the blockade of *MYC* dosage compensation in cancer with *MYC* amplification is more sensitive to the inhibition of those miRNAs *in silico*. **B.** The experimental inhibition of those 3 miRNAs in colon cancer cells with 3 different copy numbers of *MYC* suggest a higher sensitivity to the inhibition of gene dosage compensation for colon cancer cell lines with *MYC* amplification (fluorescence+brightfield microscopy including the result of the image analysis pipeline in CellProfiler using Hoechst DNA staining (blue) to count nuclei and Propidium Iodide to stain dead cells (red)). **C.** The quantification of dead cells confirms a significant difference in cell death in cells treated with antimiRs compared to cells treated with the negative control RNA for HT29 and SW-620 cells (n=3, the differences become significant after 36 hours of incubation for HT29 cells and after 24 hours for the SW-620 cells with a p<0.01).

A sudden increase in *MYC* expression in a therapeutic context may have the potential to induce cytotoxicity, suggesting that the blockade of *MYC* dosage compensation has a promising therapeutic range that increases with the extent of dosage amplification. In order to confirm this hypothesis experimentally, we chose 3 colon cancer cell lines of the NCI60 panel with the widest variation in *MYC* copy numbers, the HCT-15 (copy number of 2), the HT-29 (copy number of 4) and the SW-620 (copy number of 7). We tested the effect of blocking *MYC* dosage compensation by transfecting the 3 cell lines with increasing concentrations (from 10 to 30 pM) of a mixture of anti-miR17, anti-miR19a and anti-miR20a and the respective concentrations of a negative RNA control. The cells were incubated and monitored by live microscopy for 72 h in the presence of Hoechst to stain all nuclei and Propidium Iodide to stain dead cells. After acquisition, images were analyzed using an image analysis pipeline programmed in CellProfiler to quantify the numbers of live and dead cells over time (Figure 7B). The qualitative observation of the images indicated no difference in cell growth and cell death between the anti-miRs and control transfected HCT-15 cells, whereas the difference between those two conditions increased slightly for the HT-29 and much more for the SW-620 cells (Figure 7B).

The quantification of cell death for several replicates showed that cell death increased specially in SW60 cells treated with 30 pM of the anti-miR mixture compared to the negative control (Figure 7C, lower panel). These results indicate that cells with higher *MYC* copy numbers are more sensitive to the blockade of gene dosage compensation by the perturbation of miR17, miR19a and miR20a. This suggests that aneuploid cancer cell lines with increased copy numbers are sensitive to blockade of gene dosage compensation.

### Breast Cancer patients with MYC dosage compensation have lower survival than patients with a non-compensated configuration

In order to further validate the relevance of gene dosage compensation in cancer and its potential as therapeutic target, we set out to investigate the emergence of *MYC* and *STAT3* dosage compensation in the breast cancer dataset of the Tumor Cell Genome Atlas (TCGA). First, we tried to fit the minimal model of *MYC*-*STAT3* dosage compensation to all of the data using COPASI but the objective function was very high (not shown). We noticed a huge heterogeneity in the objective function values of the individual patient cases, suggesting indeed a heterogeneous configuration in this network. Therefore, we used the model as a tool to characterize the patient cases by fitting the model to the data of each individual experiment (case) by automatically constructing patient-tailored models to obtain a set of parameters for each of them. Next, we performed individual parameter scans to further characterize the cases in function of their response to copy number increases for each of the candidate genes for which a minimal compensation model was postulated (*MYC*, *STAT3*, *STAT5A* and *STAT5B*). Therefore, we increased the gene copy number from 1 to 5 and observed the increase in gene expression: a sub-linear increase denotes the capability of the respective network configuration to achieve dosage compensation. We observed a large heterogeneity in *MYC* and *STAT5A* dosage compensation and no dosage compensation capability was observed for *STAT3* with only few cases for *STAT5B* (Figure 8A), suggesting that the configuration of gene dosage compensation circuits may vary across patients.

**Figure 8.**
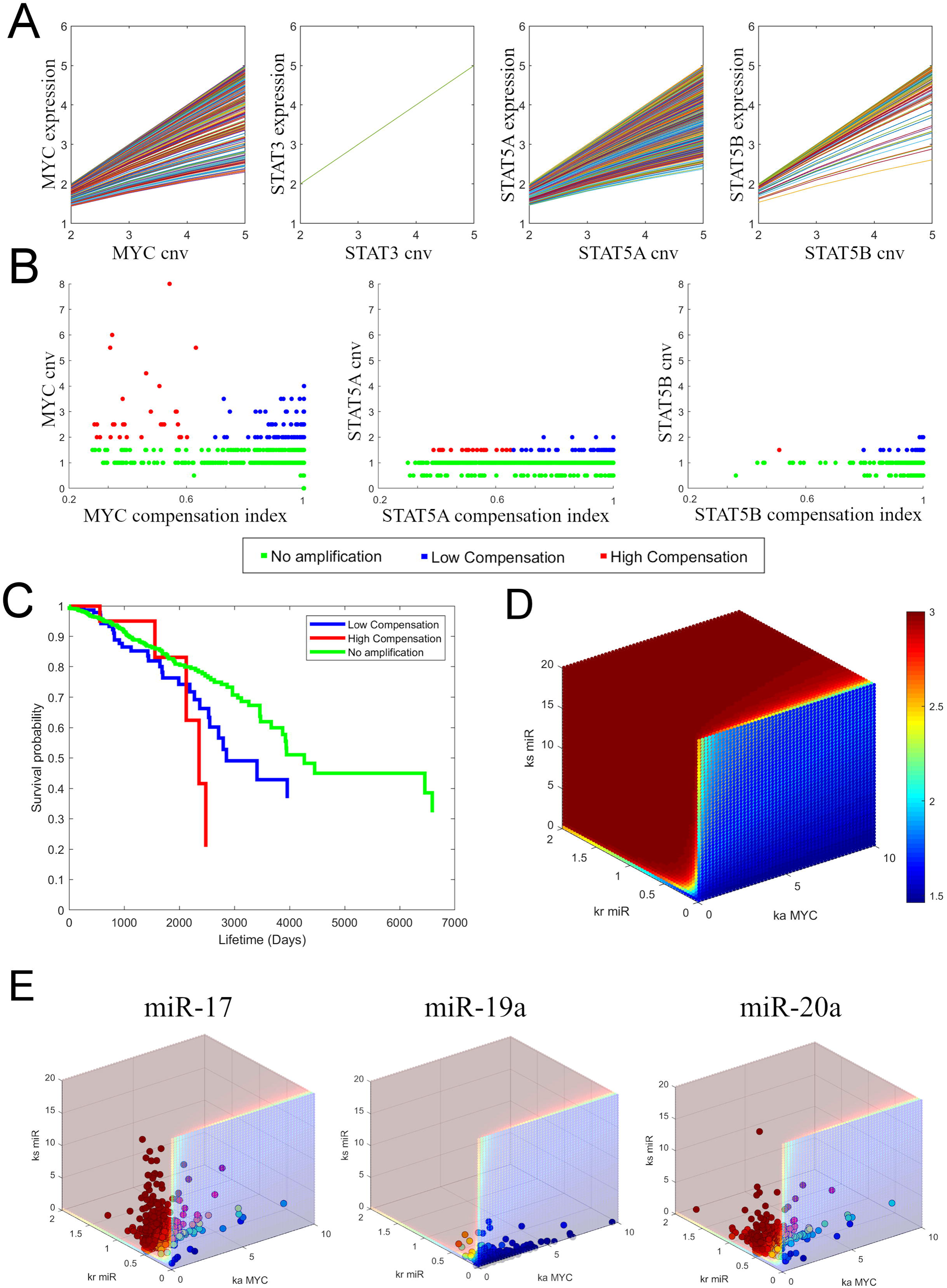
Breast Cancer patients show heterogeneity in gene dosage compensation leading to differences in patient survival. **A.** Assessment of gene dosage compensation by individual parameter scans of the gene copy numbers of *MYC*, *STAT3*, *STAT5A* and *STAT5B* for each patient-tailored model individually fitted, showing varying degrees of heterogeneity in gene dosage compensation. **B.** Patient case separation by the amplifications (experimental CNV) and compensation behavior (compensation index *in silico*) for *MYC*, *STAT5A* and *STAT5B*. **C.** Survival probabilities for the patient cases divided according to the amplification and compensation behavior of *MYC*. **D.** The compensation landscape of model parameters for the models of the TCGA breast cancer patients calculated by a triple-parameter scan on the respective ranges of ks (synthesis rate of the miRNA), ka (activation parameter of the TF) and kr (repression parameter of the miRNA) under the two different copy numbers of *MYC* (1 and 3). The ratio between these two scans represents the three-dimensional landscape of gene dosage compensation for the range of parameters obtained by the individual fitting of the patient-tailored models, where the blue region represents the space of parameter values leading to compensation and the red region the one without dosage compensation. **E.** The parameter sets of the individual miRNA-mediated regulatory motifs are interpolated within this landscape indicating that miR-19a is responsible for gene dosage compensation for most models of *MYC* dosage compensation in breast cancer patients.

The response of the corresponding network configuration also sheds light on the importance of gene dosage compensation in cancer. We first classified the patients according to their gene amplification and compensation capability of their fitted models (see STAR methods: material and methods: Calculation of Compensation Index). The lower the compensation index, the higher the capability of the corresponding network configuration to perform compensation. As for *STAT3* in CCLE, there were very few cases of amplification for *STAT5A* and *STAT5B* to evaluate the effect of gene dosage-compensation on patient survival but *MYC* amplifications were very frequent. We could observe that the higher the extent of this amplification the higher the separation of cases in terms of *MYC* compensation leading to a clear separation of 2 groups within the amplified cases (Figure 8B). We therefore separated the cases in three groups: “No amplification” (CNV lower or equal to 1.5), and for those with higher amplification a group with “low compensation” (*MYC* expression higher than 3.7) and a group with “high compensation” (*MYC* expression lower than 3.7) (Figure 8B). We plotted their survival curves implementing a Kaplan-Meier plot in Matlab (ecdf function, MathWorks) to compare the survival probabilities among the groups (Figure 8C). The significance in their differences was assessed by a Cox proportional hazards regression (coxphfit, Matlab). We observed no significant difference between the “No amplification” group and the amplified cases with “low compensation” (p = 0.1). In contrast, we observed a significant decrease in the survival probability of the patients with “high compensation” compared to those with “low compensation” (p = 0.009) and compared to the cases of “No amplification” (p = 0.007). This result suggests that *MYC* dosage compensation enables cancer progression towards malignancy and it could indeed represent an important therapeutic target to prevent this progression.

Finally, to further characterize the nature of *MYC* dosage compensation in these models of breast cancer patients, we defined the new ranges of the 3 parameters previously identified to best partition the parameter landscape into the non-compensated and compensated spaces (Figure 5E). Using these ranges, we performed the corresponding triple parameter scan and could identify again both regions of compensation behavior for the breast cancer cases (Figure 8D). Using the values of the fitted parameters for each patient, we used the N-D nearest point search algorithm (dsearchhn, Matlab, Mathworks) to obtain the compensation value for each patient for that particular miRNA-mediated feedback motif. We observed that miR-17 and miR-20a mediate *MYC* dosage compensation only in a small fraction of cases whereas most cases of breast cancer patients are compensated by the feedback motif regulated by miR-19. This suggests that miR-19 inhibition would represent a potential therapeutic target to block *MYC* dosage compensation in breast cancer.

## Discussion

We report in this work a systems biology approach which successfully allowed us to test and verify the hypothesis we made that gene dosage compensation in cancer is mediated by the emerging properties of complex miRNA-TF networks, which control the expression of key genes with altered copy number.

In endogenous transcription networks, interactions of miRNAs and transcription factors have been reported to assemble complex motifs including negative feedback loops, positive feedback loops, coherent feedforward loops, incoherent feedforward loops, miRNA clusters and target hubs leading to non-linear, systems-level properties such as bistability, ultrasensitivity and oscillations (Lai et al., 2013; Vera et al., 2013). Given this complexity, gene dosage compensation could arise as an emergent property of the system, meaning that a differential expression analysis of gene targets or miRNAs would be insufficient to identify the underlying mechanism of gene dosage compensation. Classical approaches in miRNA research start with the identification of dysregulated miRNAs related to disease and require extensive molecular and cellular biology work to validate the gene targets related to the phenotype of interest. This is inefficient because each miRNA can alter the expression of hundreds of genes by only 1.5- to 4-fold (Vera et al., 2013) and it is the cooperative or synergistic effect of miRNA networks that makes them robust regulators (Herranz and Cohen, 2010; Matsuo et al., 2010). Thus, the identification of critical miRNAs involved in gene-dosage compensation networks requires the analysis of a dynamical model of those interactions by means of sensitivity analysis combined with predictive simulations to identify processes that can become potential therapeutic targets (Lai et al., 2013; Vera et al., 2013).

To achieve this, we developed a biocomputational platform to identify the critical miRNA-mediated gene regulation and to perform simulations of the systems-level effects of perturbations in miRNA networks related to disease. This is accomplished by first casting a wide net and generating a large network out of a database of 450,000 experimentally validated regulatory interactions, which is translated to a dynamical model. The process then greatly reduces this model, based on biology inspired *in silico* experiments and sensitivity analysis, to arrive at a small model that translates the critical regulatory mechanism for gene dosage compensation (only 11 interactions). Due to the redundancy in gene dosage compensation, it would be unfeasible to identify such mechanisms by single or even double inhibitions using a functional genomics approach alone. To our knowledge, this is the first example of such a top-down approach in miRNA research which success relies on network simplification based on sensitivity analysis and biology inspired-experiments *in silico*.

Our current approach, in spite of some limitations, facilitated the analysis of potential mechanisms of gene dosage compensation in a variety of aneuploid cancer cells. Our data indicate that these cells, whatever their cancer origin, regulate dosage compensation at the transcriptional level. To our knowledge, there is still few evidence for regulation at that level, specifically in mammalian cells except for Down Syndrome (Kahlem et al., 2004; Lyle et al., 2004) and the case of sex chromosomes inactivation. Three major mechanisms have been studied for sex chromosomes: inactivation of one of the X chromosomes in female embryos in mammals (Heard et al., 1997); doubling of transcription of genes on the single male X chromosome in *D. melanogaster* (Lucchesi and Kuroda, 2015) and halving the expression of genes on the two X chromosomes in hermaphrodites of *Caenorhabditis elegans* (Meyer, 2005; Meyer et al., 2004). Although some of these mechanisms have been related to epigenetic modifications and other nonconding RNAs (Park and Kuroda, 2001), there was no evidence of a role of miRNAs in specific gene dosage compensation.

In this work, we report another mechanism mediated by the emerging properties of complex miRNA-TF regulatory networks. Its structural analysis revealed that only those with TF function can be subjected to gene dosage compensation. Once we enriched the network topology for sensor loops of experimentally validated interactions to simplify the regulatory network, we were able to model this mechanism and validated the hypothesis for the genes *MYC* and *STAT3*. A deeper analysis of this network enabled us to understand the underlying mechanism of gene dosage compensation by the reduction of model complexity into a minimal model that is still able to display the same behavior of dosage compensation for both genes. Indeed, an important characteristic in this type of biological networks is its capability to adapt to fluctuating concentrations of its biomolecular components. The mechanism includes coherent and incoherent feed forward loops (FFL) and feedback loops (FBL) that help the network to adapt from such fluctuations (Carignano et al., 2019; Osella et al., 2011). Moreover, it has been shown, using synthetic transcriptional and post-transcriptional incoherent feed-forward loops that a gene product can adapt to changes in DNA template abundance, supporting a previously hypothesized endogenous role in gene dosage compensation for such motifs (Bleris et al., 2011; Shimoga et al., 2014).

The reduced model has a higher explanatory power than the original (large) network, which enabled us to design a strategy for experimental validation using a genetic tug-of-war approach. Under the pressure of the dosage compensation circuits, the exogenous *MYC* overexpression leads to the down-regulation of endogenous *MYC*, confirming thereby the ability of these circuits to perform gene dosage compensation and a possible involvement of the 3 miRNAs in those regulatory circuits. This novel method fills a technical gap for future experimental validation of gene dosage compensation at the transcriptional level.

Upon this experimental validation, the question emerges of whether a dosage-altered gene is critical, or whether it require compensation to maintain cell viability. The gene *MYC* (*MYC* proto-oncogene, bHLH transcription factor), aka c-*MYC*, is a well-known and studied transcription factor involved in proliferation, cell growth, cell differentiation and apoptosis. It is estimated that *MYC* regulates 15% of all genes. It is as well an oncogene, active in 70% of cancers, and overexpressed in various types of cancer (Dang, 1999). *MYC* however is also reported to act as tumor suppressor in leukemia (Uribesalgo et al., 2012). Whereas *MYC* dysregulation may lead to cancer it also triggers cell suicide (Nilsson and Cleveland, 2003). Therefore, it is not surprising that (exo)*MYC* expression in our experimental setup led to cell death, supporting the prediction of a study of gene dosage compensation in aneuploid yeast, which suggested that the phenomenon of dosage compensation occurs at genes which are most toxic when overexpressed (Hose et al., 2015).

Those results also suggest that the inhibition of the three miRNAs compensating *MYC* could induce cell death. The miR17, miR19a and miR20a are actually all co-regulated like the miRNA cluster miR-17-92. This is considered an oncogenic cluster and is actually called OncomiR-1 since it is overexpressed in several types of cancer (for review see (Fuziwara and Kimura, 2015))). Several reports showed that these miRNAs form important network motifs with *MYC* in B-cell lymphoma (Mihailovich et al., 2015), with *E2F/MYC* (Li et al., 2011) and with *STAT3* in retinoblastoma (Jo et al., 2014). Interestingly, using our model, the *in silico* inhibition of those 3 miRNAs led to a dosage-dependent increase in *MYC* expression, while the *in vitro* inhibition of those miRNAs led to a dosage-dependent increase in cytotoxicity, suggesting that dosage-altered cancers are more fragile to blockade of the dosage compensation mechanism.

Based on our observations and computational simulations, we suggest that cancer has a robust Achilles-heel due to an increased sensitivity to perturbations in these circuits, which is not necessarily reflected in differences in individual miRNA expression levels but rather at more complex systems-level properties. The identification of targets that may block dosage compensation could lead to a strategy where the overexpression of these genes, and others under their influence, could exploit this fragility of cancer cells. This strategy is promising inasmuch the overexpression of these important transcription factors seem to be more sensitive to blockade of gene dosage compensation when their copy numbers are more amplified. Indeed, the identification of miRNAs controlling cancer robustness has a strong therapeutic potential since miRNAs are becoming attractive targets for therapy, as shown for the first time by miRNA-122 against hepatitis C infection and hepatic cancer (Lindow and Kauppinen, 2012) and for, at least, another 7 miRNA mimics/inhibitors being tested in clinical trials (Rupaimoole and Slack, 2017).

Nevertheless, the potential therapeutic application of a strategy targeting gene-dosage compensation in cancer cells requires a systematic evaluation of the extent of the copy number alterations and their compensation across large data sets, such as CCLE and TCGA, for the identification of further candidate genes, the availability of more data to obtain more accurate models, and the evaluation of the tumor heterogeneity in gene dosage compensation. The further analysis of the current models of gene dosage compensation or even other genes could reveal novel pan-cancer or cancer type-specific targets. Our analysis of TCGA data for samples of breast cancer patients revealed that these tumors have differential configurations in these compensation circuits as we observed different degrees of *MYC* dosage compensation and a negative impact on survival of patients with a higher compensation index. These findings suggest that *MYC* dosage compensation contributes to a status of pro-tumoral stability enabling cancer cells to evolve into more malignant phenotypes. It has been proposed that between cell cycling and apoptosis there is a range of *MYC* (and/or E2F) activity associated to carcinogenesis and referred to as the cancer zone (Aguda, 2013). The future determination of the molecular determinants of compensation (e.g. mutations in interacting sequences of miRNA-targets) could lead to the identification of biomarkers to direct precision medicine strategies against gene dosage compensation to induce apoptosis out of that cancer zone.

In conclusion, the present work led to the construction of a mathematical model of gene dosage compensation and formulated a model-driven hypothesis for the identification of novel targets against aneuploid cancer. We developed a computational platform including bioinformatics and dynamical modeling tools for any application in miRNA research (BioNetUCR, available here: <u>bionet.prislab.org/download</u>. In addition, we presented a rationale to identify dosage compensation circuits and a novel genetic tug-of-war approach to experimentally validate those regulatory motifs. Besides, we evaluated the therapeutic potential of MYC dosage compensation to target aneuploid cancer. Future work is required to confirm the effect of gene dosage compensation on patient survival and to identify other compensated genes to direct personalized precision therapies against cancer. Altogether, the current results contribute to the identification of a stability core of essential genes, which manipulation of specific nodes has the potential to become a novel approach to specifically target aneuploid cancer cells.

### Limitations of the study

- The current criterion to identify candidate genes under dosage compensation does not include a direct metric of the number of amplifications of the gene of interest since it is only based on standard deviations.
- We could only describe gene dosage compensation circuits for a small fraction of the candidates, suggesting that other mechanisms of compensation might be at work or there is an indirect effect of the dosage compensation of one gene impacting on others, which we did not study in this work.
- The NCI60 dataset is limited to identify candidates and the CCLE data is very heterogeneous to fit large-scale models of miRNA-TF interactions. CCLE or TCGA tumor-type specific models may be a better option for future studies.
- There is heterogeneity in the parameters of individual cell lines probably due to mutations in the TF-miRNA-target interaction sequences or other epigenetic mechanisms that are not included in our models. Therefore, it is not possible to determine a specific set of model parameters of the compensation models but a parameter landscape of gene dosage compensation.
- The construction of the regulatory networks depends on the availability of experimentally reported interactions. A methodology is required to prioritize which putative interactions should be included.
- Our genetic tug-of-war techniques implies extensive molecular and cellular biology work to validate the dosage compensation for only one gene at the time.
- The pre-clinical validation of these findings requires the development of miRNA sponges for multiple miRNAs since gene dosage compensation seems to be highly redundant in cancer.

## Acknowledgments

We thank the initial contribution of Dr. Michael Kesling for his advises on the project formulation and the data analysis. This project was funded by the grant FEES-CONARE (Costa Rica), University of Costa Rica and Universidad Estatal a Distancia (UNED). Rodrigo Mora was supported by the Georg Foster Grant of the Alexander von Humboldt foundation (Germany). Carsten Geiß was supported by the FAZIT-STIFTUNG. Support by the IMB Flow Cytometry Core Facility (Mainz, Germany) is gratefully acknowledged. PM thanks the NIH for supporting the development, use and dissemination of COPASI (grants GM080219 and GM137787)

## Author contributions

Conceptualization MA, ARV, PM and RMR. Methodology MA, CG, JLAA, DBE, GO, LARM, EB, GVV, YOO and RMR. Software MA, JTC, GO, LARM, EB, ASC, FSC. Validation JGC, FSC, SQB, ARV and PM. Formal Analysis AM, CG, ARV, PM and RMR. Investigation MA, CG, DBE, JLAA, JGC, ASC and RMR. Resources, ARV, ASC, FSC, PM and RMR. Data Curation, AM, JTC, GO, EB and RMR. Writing-Original Draft, AM, CG and RMR. Writing-Review&Editing, JGC, SQB, ARV, PM and RMR. Visualization, FSC, JTC, LARM and RMR. Supervision SQB, ARV, PM and RMR. Project Administration ARV, PM and RMR. Funding Acquisition ARV, PM and RMR.

## FIGURE LEGENDS

**Figure S1.**
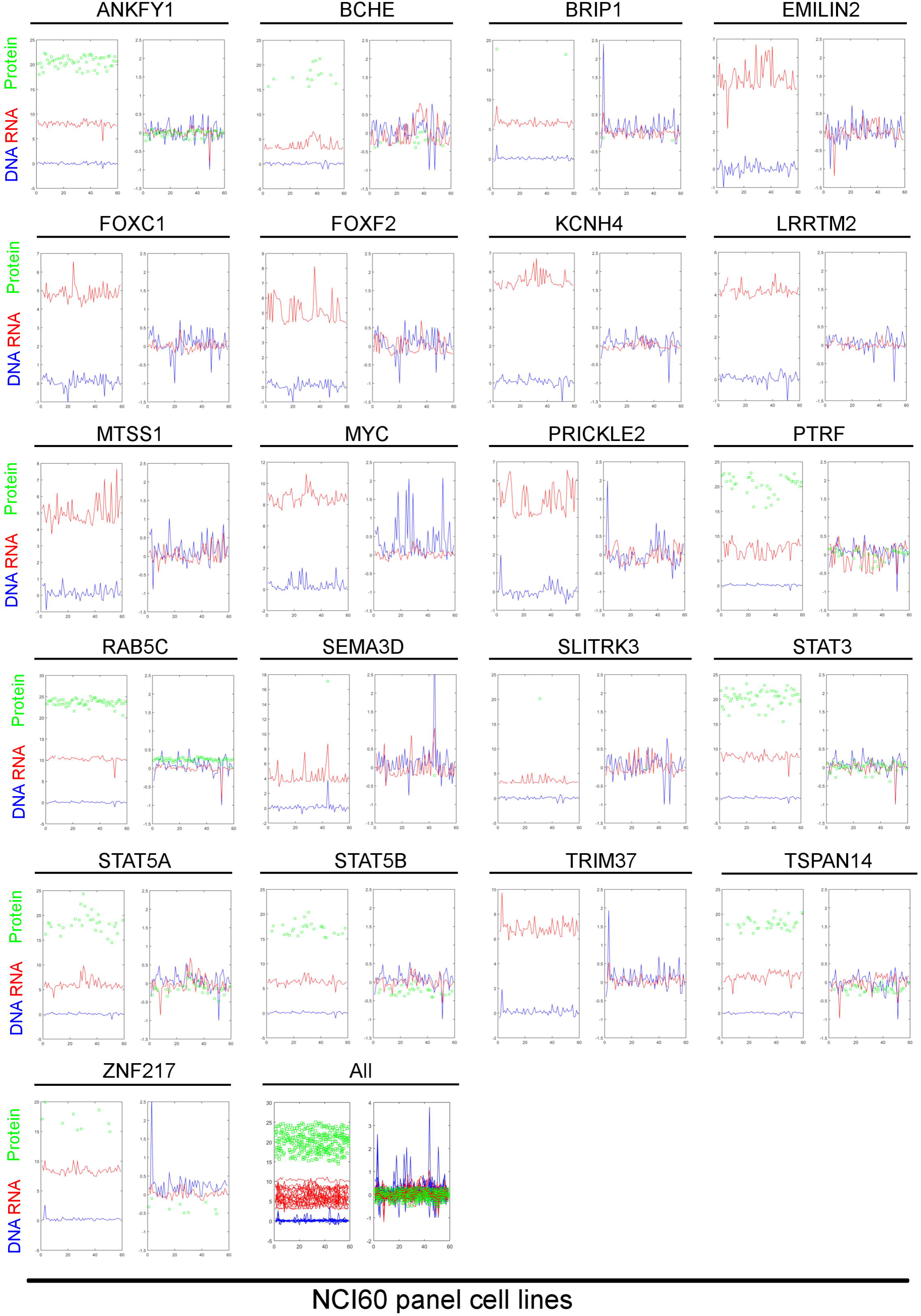
DNA, RNA and protein levels for all candidate genes. (DNA, blue), gene expression (RNA, red) and protein levels (protein, green) across the NCI60 panel. The absolute values are shown on the left panel across the 59 cell lines and the right panel corresponds to the log2 values normalized to the averaged RNA and protein of the diploid cell lines for the respective gene (Normalized to diploid).

**Figure S2.**
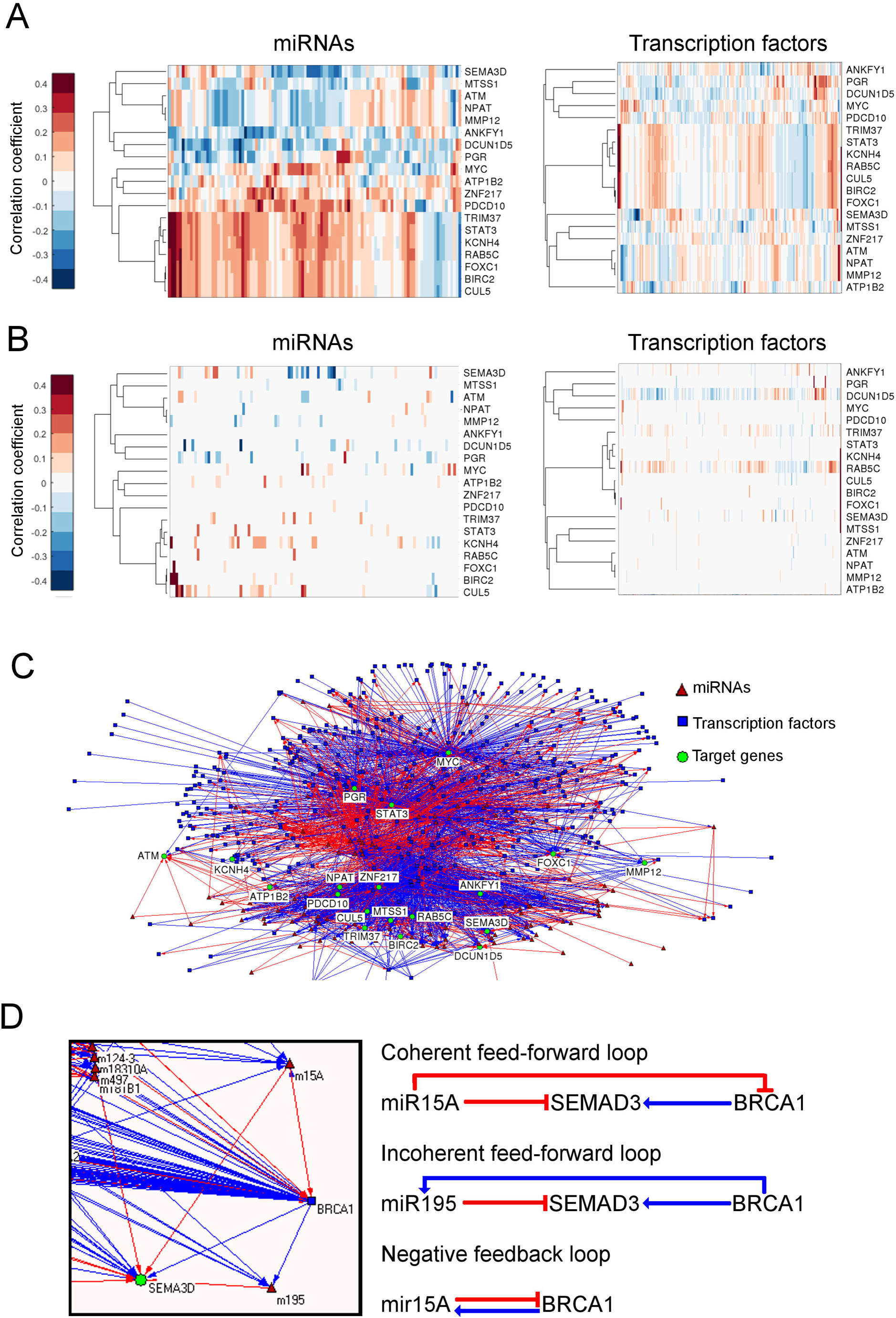
Putative miRNA-transcription factor interaction network linking all candidate target genes. **A.** Correlation coefficients of copy number variations of the candidate genes and miRNA/transcription factor expression across the NCI60 panel. **B.** Correlation coefficients of copy number variations of the candidate genes and miRNA/transcription factor expression across the NCI60 panel, only for those putative interactions present on the network in C. **C.** Network of putative interactions among target genes, miRNAs and transcription factors (red indicates repression and blue activation) including different types of regulatory loops formed by target genes, miRNAs and TFs. The Circos representation includes the chromosomal distributions of the different genes of the model (1-22). **D.** Example miRNA-TF regulatory circuits including negative feedback loops, coherent feed-forward loops and incoherent feed-forward loops.

**Figure S3.**
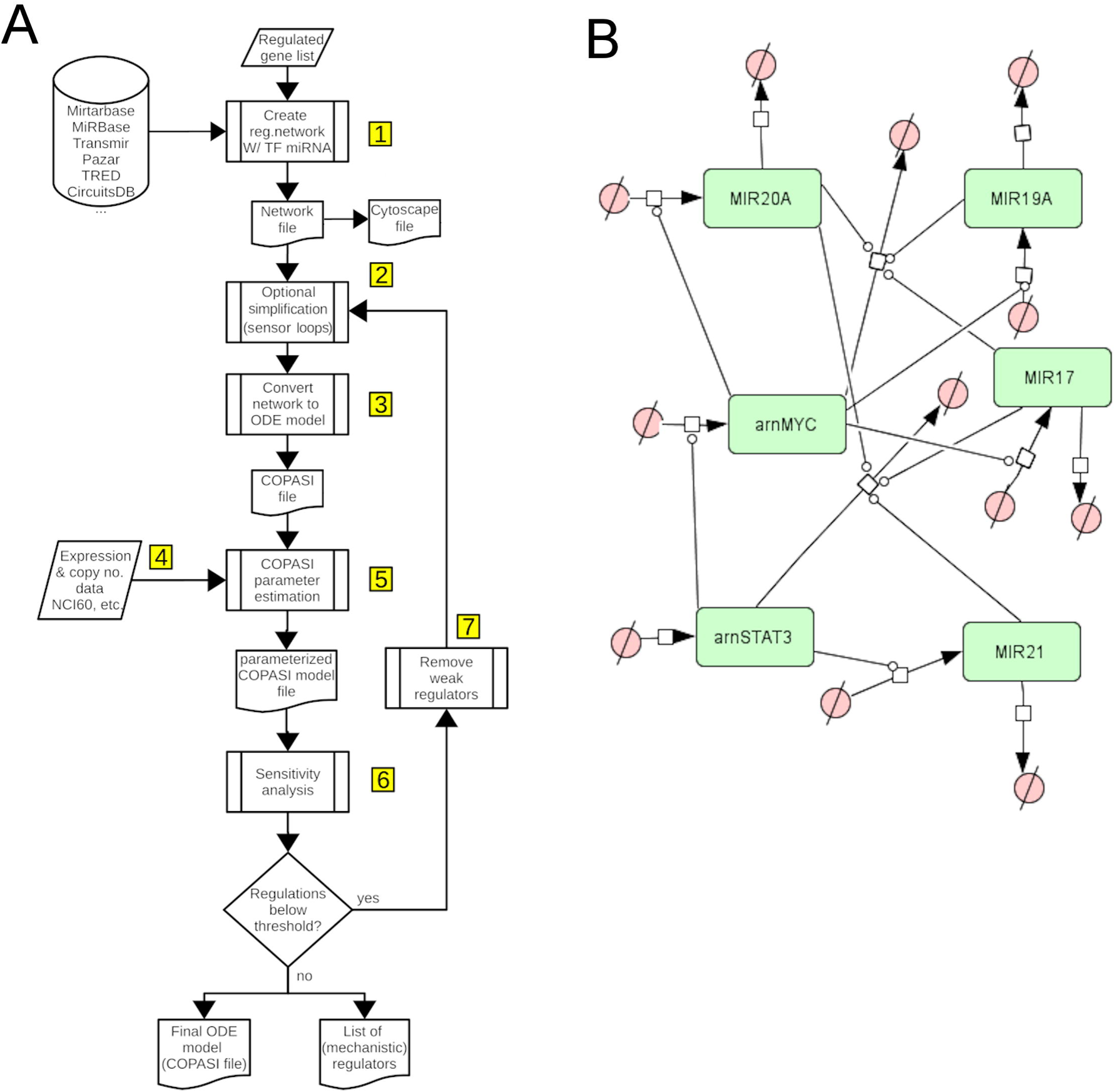
Flowchart of the biocomputational platform for the step-wise construction of mathematical models of miRNA-TF interactions. **A.** The input is a list of genes of interest for the platform to automatically construct a network topology that can also be visualized using Cytoscape (step 1). Optional simplification steps are included to remove or add nodes or interactions and to enrich the model with sensor loops, this is the only manual step to configure the network (step 2). The network is automatically converted to an ODE mathematical model for COPASI (step 3) and integrates experimental data including expression and copy numbers of NCI60, CLLE or TCGA (breast cancer) or any kind of data provided. This is automatically incorporated in the definition of the parameter estimation task of COPASI to obtain a model ready for fitting (step 5). This is done in COPASI together with a statistical evaluation for goodness-of-fit and sensitivity analysis to identify the nodes with lowest fitting and lowest control on the genes of interest as candidates to be removed in each iteration (step 7) to generate a new network and mathematical model to be fitted until there is no significant difference between the experimental and simulated data. The fitted model can now be used to perform simulations, sensitivity and stability analysis in COPASI to investigate which are the main regulators for the phenotype of interest obtaining minimal models of miRNA-TF regulation. **B.** SBGN representation of the minimal model of MYC-STAT3 regulation. State transition arrows indicate synthesis or degradation, catalysis lines (circular-head) indicate activation and flat-head-lines indicate inhibition.

**Figure S4.**
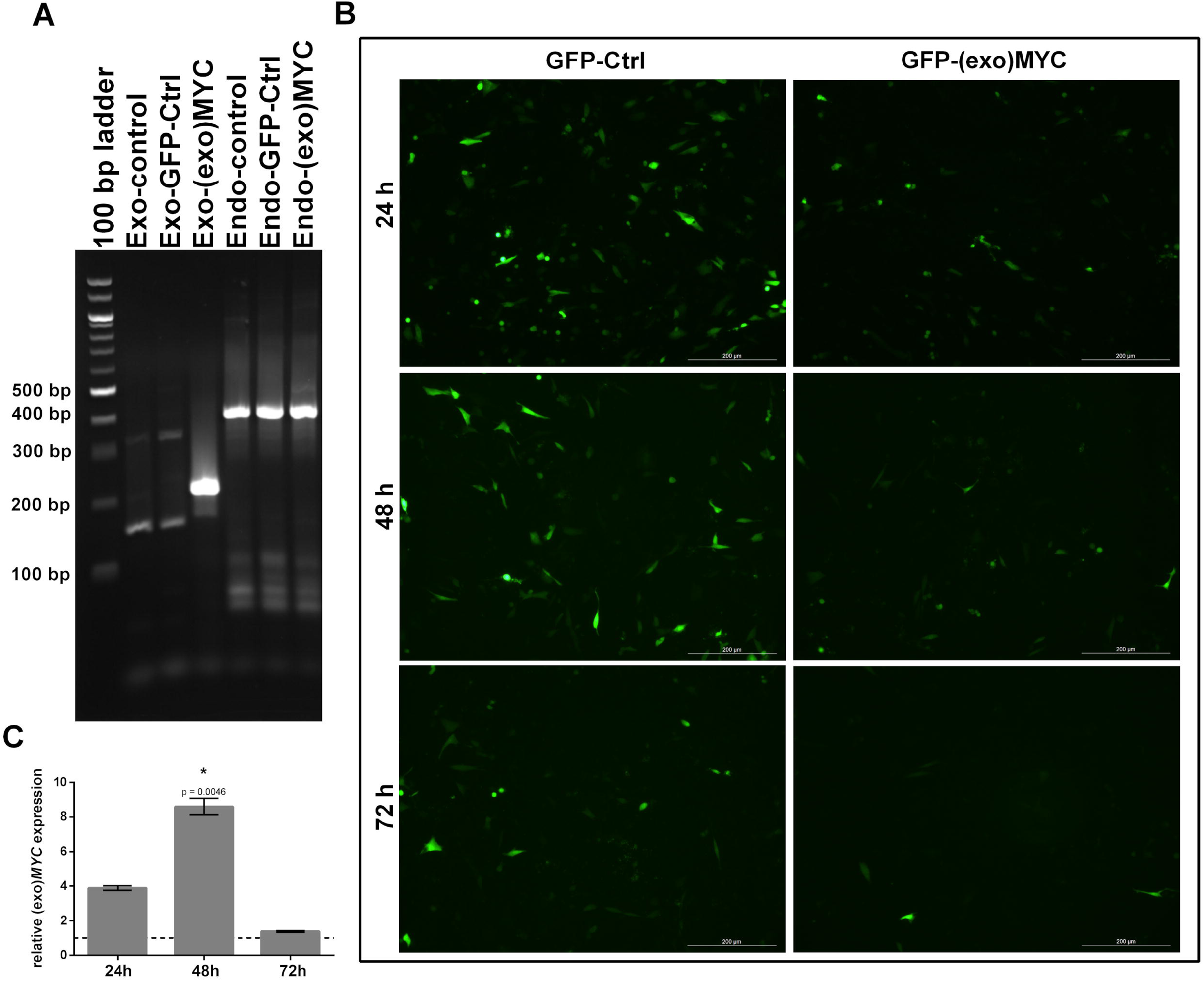
– Characterization of the human GB cells overexpressing exogeneous *MYC*. **A.** Analysis on a 2% agarose gel of amplicons after quantitative RT-PCR. Expected product sizes were 216 bp (exogeneous *MYC*) and 404 bp (endogenous *MYC*). Lane 1: 100 bp ladder; lanes 2-4: untransfected, GFP-Ctrl transfected and GFP-(exo)*MYC* transfected NCH82 cDNAs analyzed using exogenous *MYC* primers; lanes 5-7: untransfected, GFP-Ctrl transfected and GFP-(exo)*MYC* transfected NCH82 cDNAs analyzed using endogenous *MYC* primers. **B.** Representative microscopic images of transfected NCH82 cells 24, 48 and 72 h after transfection. Left panel: cells transfected with the GFP-Ctrl plasmid; right panel: cells transfected with the GFP-(exo)*MYC* plasmid. Scale bars: 200 µm. **C.** Quantitative RT-PCR detection of (exo)*MYC* expression in the *MYC* positive population relative to the *MYC* negative population after sorting of GFP-(exo)*MYC* transfected cells. (n = 3); t-tests were performed to analyze means of samples with a significance defined by an α of 0.05. The resulting p-values were corrected for multiple comparisons using the Holm-Sidak method.

## STAR Methods

### RESOURCE AVAILABILITY

#### Lead contact

Further information and requests for resources, datasets and programing code should be directed to and will be fulfilled by the lead contact, Rodrigo Mora-Rodríguez.

#### Materials availability

The plasmids generated in this study for the experimental demonstration of *MYC* dosage compensation by a genetic tug-of-war pSBbi-moxGFP-P2A-T2A-Puro and pSBbi-moxGFP-P2A-T2A-c*MYC*-Puro will be available in Addgene (176892 and 176892).

#### Data and code availability

The latest version of BioNetUCR, our platform for the automatic construction of large-scale networks of mathematical models can be found here: <u>bionet.prislab.org/download</u> and is publicly available as of the date of publication. It also includes the NCI60 data set. The CCLE data set can be found here: https://data.mendeley.com/datasets/t27jvj4vz2/draft?a=62a3d9f2-594b-46cc-8f05-592a6ad6e2bb. The mathematical models of *MYC/STAT3* dosage compensation are included as supplementary files. MATLAB programming codes for data analysis, specific tasks or plotting is available upon request. This paper analyzes existing, publicly available data. Any additional information required to reanalyze the data reported in this paper is available from the lead contact upon request.

### EXPERIMENTAL MODEL AND SUBJECT DETAILS

#### Human glioblastoma cell lines

The human primary glioblastoma cell line NCH82 was generated at the Department of Neurosurgery, Heidelberg University Hospital (Heidelberg, Germany) (Karcher et al., 2006). Cells were cultured in cDMEM [DMEM (Sigma-Aldrich, St. Louis, MO, USA), 10% heat-inactivated FCS (Sigma-Aldrich), 2 mM L-Glutamine (Gibco Invitrogen, Carlsbad, California, USA), 50 µg/ml Gentamicin (Gibco Invitrogen)] at 37°C and 5% CO_2_.

#### Human Colon Cancer Cell Lines

The human colon cancer cell lines SW-620, HCT-15 and HT-29 were obtained as part of the NCI60 collection (DTP, NIH). Cells were cultured in RPMI [GIBCO, Invitrogen, Carlsbad, California, USA), with 2 mM L-Glutamine (Gibco Invitrogen, Carlsbad, California, USA), 50 µg/ml Gentamicin (Gibco Invitrogen)] at 37°C and 5% CO_2_.

### METHOD DETAILS

#### Experimental Methods

##### Anti-miRNA transfections and cytotoxicity assays

Optimized densities of colon cancer cells lines were seeded in 96 well plates (Greiner Bio-One µclear) to evaluate the cytotoxicity of the miRNA inhibitors for 72 h: 40,000 SW-620 cells, 20,000 HCT-15 and 40,000 HT-29 cells were plated per well in 100 µL of RPMI medium supplemented with 10% of fetal serum (GIBCO), 24 h before the start of the experiment and incubated at 5% CO_2_ and 37°C. Next, cell nuclei were labeled using Hoechst 33342 (Invitrogen™ H3570, 1.25 µg/mL final concentration) for 10 minutes before washing. Afterwards, the cells were transfected with the mixture of 3 antimiRs: i. *mir*Vana^®^ miRNA inhibitor, hsa-miR-17-5p-assayID: MH12412 (ThermoFisher Scientific 4464084), ii. *mir*Vana^®^ miRNA inhibitor, hsa-miR-19a-3p-assayID: MH10649 (ThermoFisher Scientific 4464084) and iii. *mir*Vana^®^ miRNA inhibitor, hsa-miR-20a-5p – assay ID: MH10057, (ThermoFisher Scientific 4464084) or the negative control (*mir*Vana^®^ miRNA inhibitor, Negative Control #1, ThermoFisher Scientific 4464076) using Lipofectamine™ RNAiMAX Transfection Reagent (ThermoFisher Scientific 13778075) and following the Lipofectamine® RNAiMAX Transfection Protocol of the manufacturer at final concentrations of 10, 20 and 30 pmol/well. Briefly, a 1:1 mixture of lipofectamine and miRNA mixture was prepared, incubated 5 min at room temperature and 10 µL were added to each well. The cells were incubated for 6 h at 37°C with 5% CO_2_ before replacing the media. Propidium iodide (PI, Invitrogen™ P3566, final concentration of 10 µg/mL) was used to stain dead cells in a further 72 h incubation inside the imaging chamber of a Cytation 5 Imaging System (Biotek) with an imaging frequency of every 12 hours. Control wells of untransfected cells were included for each cell line. Images in the red (PI, dead) and blue (Hoechst, nuclei of all cells) channels were taken. With this information, live cell percentage was determined using the Cell Profiler image analysis software, through an image analysis pipeline designed to count total and dead cells (see supplementary pipeline file attached). The percentage of cytotoxicity was calculated by dividing PI positive cells by the total number of cells in each well.

##### Molecular design and cloning

A construct encoding the moxGFP gene (Costantini et al., 2015) followed by 2A sequences from Porcine Teschovirus-1 (P2A) and <u>Thosea asigna</u> virus (T2A) was commercially synthetized with human/mouse codon optimization (GenScript, Piscataway, NJ) and inserted into the NcoI and XbaI sites of the transposon-based plasmid pSBbi-Pur (a gift from Eric Kowarz, Addgene plasmid #60523) (Kowarz et al., 2015), to produce the vector pSBbi-moxGFP-P2A-T2A-Puro. Next, nucleotides 409-1728 of the c-*MYC* cDNA (GenBank reference sequence NM_002467.6) were commercially synthetized with human/mouse codon optimization (GenScript) and inserted into the SmaI and XbaI sites of the plasmid pSBbi-moxGFP-P2A-T2A-Puro, to generate the polycistronic vector pSBbi-moxGFP-P2A-T2A-c*MYC*-Puro.

##### Plasmid transfection in human glioblastoma cell lines

One day prior transfection, 100,000 NCH82 cells/well were seeded in cDMEM in 24 well tissue culture vessels (Greiner Bio-One, Frickenhausen, Germany) and incubated for 24 h (37°C, 5% CO_2_). At the day of transfection, old medium was replaced by 500 µl fresh cDMEM; 1 µg plasmid and 3 µl Lipofectamine™ 2000 Transfection Reagent (ThermoFisher Scientific, Waltham, MA, USA) were mixed in 100 µl Opti-MEM™ I Reduced Serum Medium (Gibco Invitrogen) and incubated for 5 min at RT. Afterwards transfection mix was added to the wells and cells were incubated for 24-72 h. Cell growth and GFP expression were documented using a Leica DM IL LED (Leica Microsystems GmbH, Wetzlar, Germany).

##### Flow Cytometry and Fluorescent Activated Cell Sorting (FACS)

Prior to analysis, cells were detached using 0.25% Trypsin-EDTA (Life Technologies, Carlsbad, California, USA), centrifuged (400xg, 5 min, 4°C) and washed with ice-cold PBS (Gibco) and stained for 15 min in PBS containing 1µg/ml propidium iodide (PI; Carl Roth, Karlsruhe, Germany). Samples for flow cytometry were analyzed using an Attune NxT Flow Cytometer (ThermoFisher Scientific). For FACS, samples of three technical replicates were pooled and analyzed using a BD FACSAria™ III Cell Sorter (BD Biosciences, San Jose, CA, USA) at the Flow Cytometry Core Facility of the Institute of Molecular Biology, Mainz. PI positive cells were discarded whereas PI negative cells were divided in a GFP positive and a GFP negative population. Cells were collected in QIAzol Lysis Reagent (Qiagen, Hilden, Germany) and stored at −80°C until total RNA extraction.

##### Total RNA isolation, cDNA transcription, *MYC* and microRNA expression analysis

After FACS, total RNA isolation was performed on the sorted cells using the miRNeasy Mini Kit (Qiagen) according to manufacturer’s instructions. RNA concentration and quality were determined using a Nanodrop 2200 (ThermoFisher Scientific). For cDNA transcription of mRNA, the FastGene Scriptase II - Ready Mix (NIPPON Genetics Europe, Düren, Germany) was used as described in the manufacturer’s protocol. For miRNA transcription, 10X M-MuLV-buffer (New England Biolabs [NEB], Ipswich, MA, USA), 1 mM adenosine triphosphate (Life Technologies), 1 mM dNTP-Mix (NEB), 10 U/µl moloney murine leukaemia virus reverse transcriptase (NEB), 0.1 U/µl poly-A polymerase (NEB), 0.05 pM cel-mir-39 as spike-in control and 0.5 µM of reverse transcriptase primers (Table 2) were added to the extracted total RNA. The reverse transcription protocol consisted of 30 min at 37°C, followed by 60 min at 42°C and a final incubation for 5 min at 65°C.

**Table 1.**
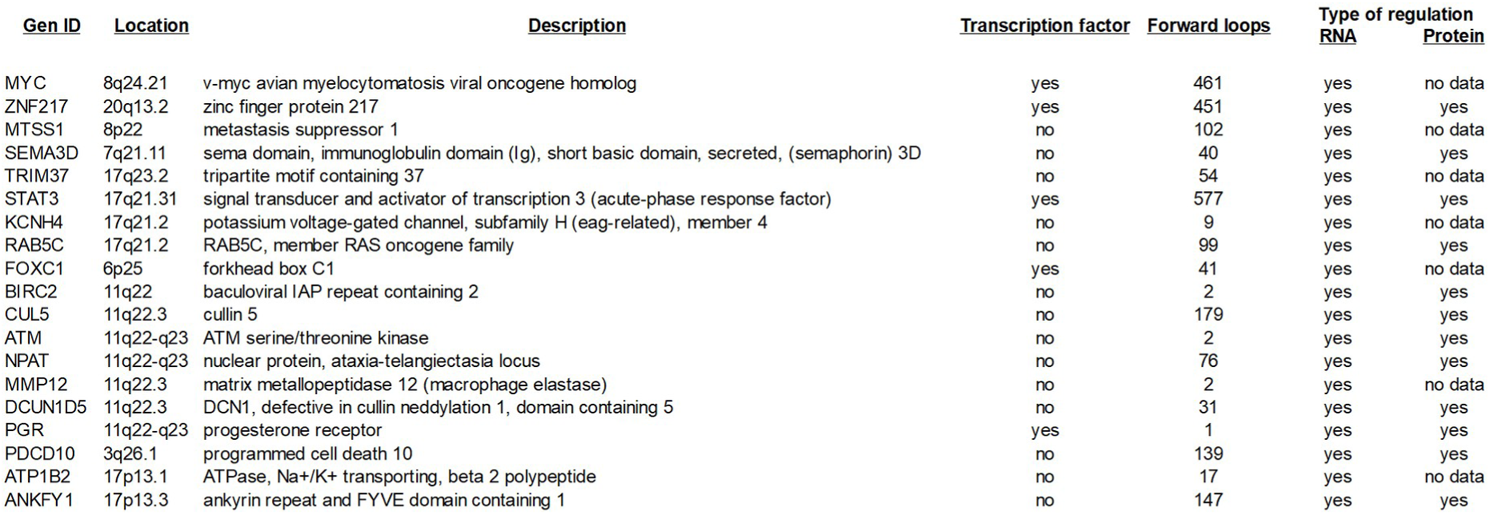
Candidate genes under dosage compensation.

| Gen ID | Location | Description | Transcription factor | Forward loops | Type of regulation |  |
| --- | --- | --- | --- | --- | --- | --- |
|  |  |  |  |  | RNA | Protein |
| MYC | 8q24.21 | v-myc avian myelocytomatosis viral oncogene homolog | yes | 461 | yes | no data |
| ZNF217 | 20q13.2 | zinc finger protein 217 | yes | 451 | yes | yes |
| MTSS1 | 8p22 | metastasis suppressor 1 | no | 102 | yes | no data |
| SEMA3D | 7q21.11 | sema domain, immunoglobulin domain (Ig), short basic domain, secreted, (semaphorin) 3D | no | 40 | yes | yes |
| TRIM37 | 17q23.2 | tripartite motif containing 37 | no | 54 | yes | no data |
| STAT3 | 17q21.31 | signal transducer and activator of transcription 3 (acute-phase response factor) | yes | 577 | yes | yes |
| KCNH4 | 17q21.2 | potassium voltage-gated channel, subfamily H (eag-related), member 4 | no | 9 | yes | no data |
| RAB5C | 17q21.2 | RAB5C, member RAS oncogene family | no | 99 | yes | yes |
| FOXC1 | 6p25 | forkhead box C1 | yes | 41 | yes | no data |
| BIRC2 | 11q22 | baculoviral IAP repeat containing 2 | no | 2 | yes | yes |
| CUL5 | 11q22.3 | cullin 5 | no | 179 | yes | yes |
| ATM | 11q22-q23 | ATM serine/threonine kinase | no | 2 | yes | yes |
| NPAT | 11q22-q23 | nuclear protein, ataxia-telangiectasia locus | no | 76 | yes | yes |
| MMP12 | 11q22.3 | matrix metalloproteinase 12 (macrophage elastase) | no | 2 | yes | no data |
| DCUN1D5 | 11q22.3 | DCN1, defective in cullin neddylation 1, domain containing 5 | no | 31 | yes | yes |
| PGR | 11q22-q23 | progesterone receptor | yes | 1 | yes | yes |
| PDCD10 | 3q26.1 | programmed cell death 10 | no | 139 | yes | yes |
| ATP1B2 | 17p13.1 | ATPase, Na<sup>+</sup>/K<sup>+</sup> transporting, beta 2 polypeptide | no | 17 | yes | no data |
| ANKFY1 | 17p13.3 | ankyrin repeat and FYVE domain containing 1 | no | 147 | yes | yes |

**Table 2:**
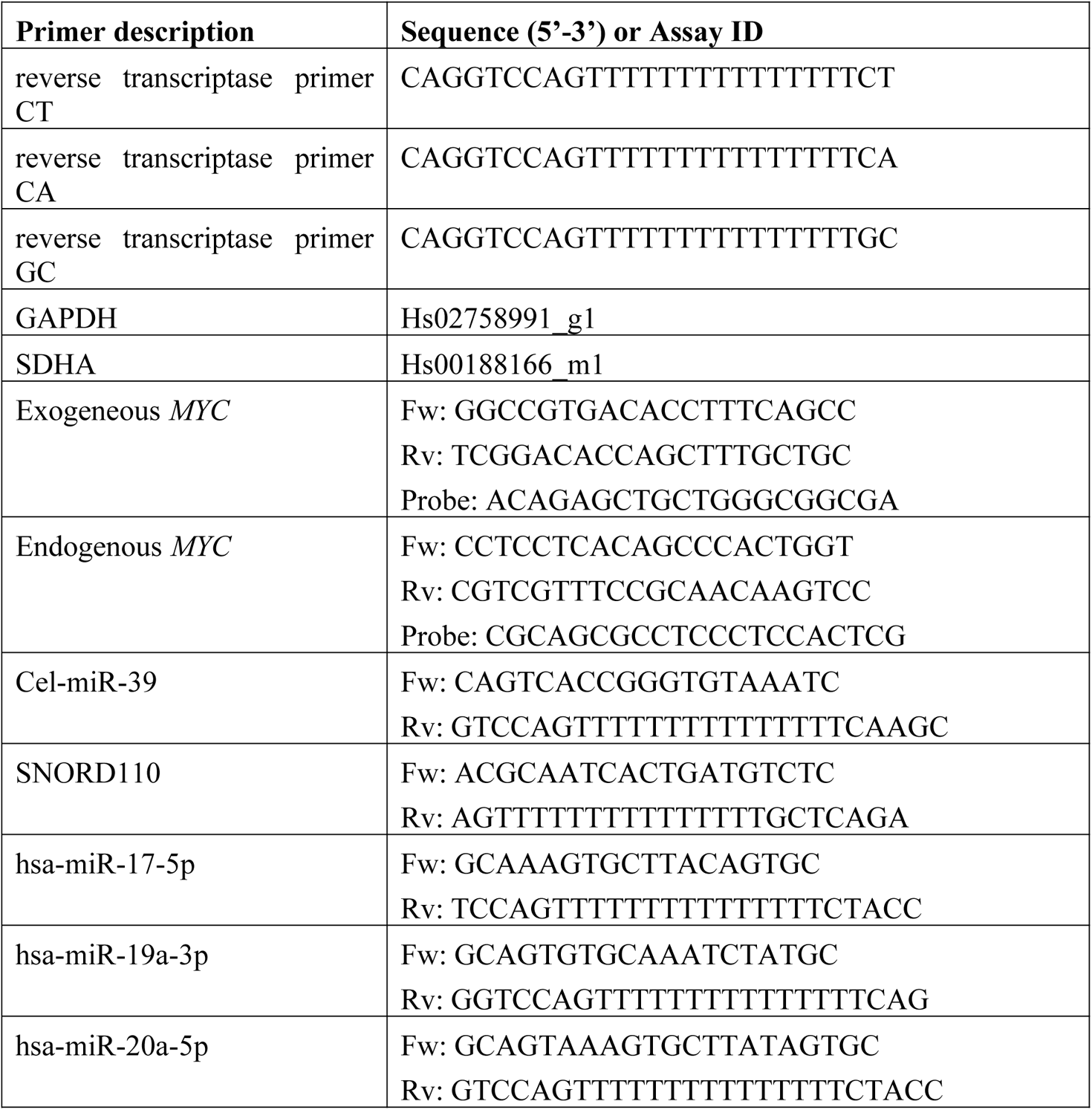
Primers and their respective sequence used for quantitative PCR. In case commercial primers were used assay ID (ThermoFisher scientific) is shown.

Quantitative PCR (qPCR) analysis was performed using 96-well qPCR plates (Sarstedt, Nümbrecht, Germany) and a StepOnePlus System (Applied Biosystems, Foster City, CA, USA). For gene expression analysis, 10 µl of qPCRBIO Probe Mix (NIPPON Genetics Europe) were mixed with 5 ng cDNA, 1 µl of the appropriate primer (Table 2) and 4 µl H_2_O. For miRNA analysis, 5 µl iTaq Universal SYBR Green Supermix (Bio-Rad, Feldkirchen, Germany) were mixed with 0.56 ng cDNA and 0.5 mM of the appropriate primer (Table 2).

Relative quantification (RQ) of gene or miRNA expression was determined using the 2^−ΔΔCt^ method (Livak and Schmittgen, 2001). *GAPDH* and *SDHA* were used as reference genes for mRNA analysis. SNORD110 was used to normalize miRNA results. Correctness of amplicons was verified using melt-curve analysis (miRNA; data not shown) and agarose gel electrophoresis (Figure S4).

Statistical analysis was performed using GraphPad Prism software. T-tests were performed to analyze means of samples with a significance defined by an α of 0.05. The resulting p-values were corrected for multiple comparisons using the Holm-Sidak method.

#### Modeling Methods

##### Data sources

Data was gathered from several sources. The primary sources were from experiments on the NCI60 panel: gene copy number (Bussey et al., 2006), RNA gene expression (Shankavaram et al., 2007) and protein expression (Gholami et al., 2013). MicroRNA related data was downloaded from Mirtarbase (Hsu et al., 2014), MirDB (Chen and Wang, 2020) and MiRBase (Kozomara et al., 2019; Kozomara and Griffiths-Jones, 2014). For background knowledge on gene regulation we relied on several sources: Transmir (Wang et al., 2010), Pazar (Portales-Casamar et al., 2009), TRED (Transcriptional Regulatory Element Database) (Jiang et al., 2007), CircuitsDB (Friard et al., 2010a)., HTRI (Bovolenta et al., 2012).

We classified the types of interactions in “experimentally validated” and “putative” according to database-specific criteria. All 407,957 miRNA-target interactions obtained from Mirtarbase were considered experimentally validated including supporting experiments such as qRT-PCR, PAR-CLIP, CLASH, HIT-CLIPS, Luciferase reporter assays, sequencing, Western blot and microarrays. MiRDB (MirTarget version 4) includes 3,375,741 predicted (putative) regulations. TransmiR v2.0 contains 3,730 literature-curated TF-miRNA regulations and 1,785,998 ChIP-seq regulations, and they were included together with Mirtarbase and TarBase experimentally validated interactions. Pazar was included to include all the transcripts of transcription factors. TRED includes 7250 curated, known and predicted regulations from transcription factor to target genes, which are graded according to promoter quality to be classified as “known, curated” (2,286), “known” (3,612) and the remaining 1,352 as predicted. According to binding quality they are classified as “known” (5,121), “Likely” (593), “Maybe” (1,314) and “predicted” (222). We classified them as putative if they have a predicted promoter quality or a predicted binding quality (6,637), or experimentally validated otherwise (216). HTRI is an open access database for experimentally validated interactions for experimentally verified human transcription factors to target genes resulting in 17,842 experimentally validated interactions between 283 transcriptions factors and 11,866 target genes. Other source of putative or predicted regulations was Harmonizome-ENCODE encompassing around 72 million functional associations between genes and proteins and their attributes such as physical relationships with other molecules. We focused on the Transcription Factor data set harvesting 1,486,116 predicted regulations between transcription factors. Finally, Circuits DB is a database of mixed miRNA/TF feed-forward regulatory circuits in human and mouse that were included as experimentally validated.

##### Gene classification using Gaussian Mixture Models

To classify genes according to their behavior, we developed a computer algorithm based on the Gaussian Mixture Model functions in MATLAB. An increasing number of components (ki) of the GMM model is added sequentially and the GMM training is performed for several iterations searching for the best fit to the experimental data. The resulting GMM is used to classify the cells of the original data set. A MANOVA test is applied to the resulting clusters to evaluate the statistical significance of adding another component to the GMM. This is done until the new component adds no further significance.

##### BioNetUCR: Software architecture development

We determined that it is possible to communicate applications developed in C# with the popular and powerful tool for network visualization called Cytoscape, using the REST style architecture. We used Visual Studio for the development. For the data storage we analyzed Neo4J and ArangoDB, both oriented to graph management. However, we ruled them out because they did not have a robust API for the integration with applications developed in C# or demanding a significant longer time for each API call in comparison to code developed directly in C#. Additionally, the installation process for the final user is simpler if no third-party data base engines are required. Therefore, the database administration was developed within the same platform creating the respective algorithms for data manipulation in RAM. Since each node has usually few arcs in comparison with the total amount of nodes, we preferred to implement the graph storage in disperse matrix. We used two vectors as the main structure. In one vector each element contains the source and target node together with the type of regulation and the second vector contains the indexes to the first vector to speed up the access. We created a class to represent the graph as designed by the user including all the genes, microRNAs and transcription factors in HUGO format. To generate the COPASI models, we used the API via DLLs. The API enabled us to create species, parameters, functions and incorporate experimental data. In addition, its usage offers greater compatibility with future COPASI versions.

##### BioNetUCR (step 1): Construction of Regulatory Network

We are interested in the regulatory network of microRNAs and transcription factors in the proximity of directly affecting the genes selected by the GMM. This list of regulated genes is input to our platform to construct a network including all the known upstream (ADD FORWARD command) and downstream (ADD BACKWARD command) regulators of these genes, both TFs and miRNAs, and the regulators of those regulators; this network was assembled from a set of regulatory interactions extracted from interaction databases (miRBase, Transmir, Pazar, Mirtarbase, TRED, CircuitsDB, HTRI) (Figure S3, step 1) to a common database of interactions. A regulatory interaction in this database is an arc which starting point is the regulator and the ending point is the target. We use a directed graph to represent this network. To construct the network, we first identify the nodes of this graph which are composed by all the regulated genes of interest as well as all the direct regulators and regulated nodes of these genes. To identify the regulators, we search in the interaction database for arcs ending in one of the selected genes. The direct regulators are all the starting nodes of each of the arcs found. Similarly, we search for arcs starting in one of the selected genes and the end node of the arc is a directly regulated gene. Hence, we have successfully identified all the nodes of the graph. Each node is a selected gene, a miRNA or TF, all of them biomolecular species relevant in the gene dosage compensation phenomenon. We then build the list of arcs of the graph. From the list of nodes or species identified, for each pair of species, if there is an arc in the interaction database linking these two species, we add this arc in the list of arcs of the graph. The lists of species and arcs identified becomes the graph representing the regulatory network affecting the selected genes of interest and they are integrated into a network file (tab separated text file of interactions). For many interactions, the database tells whether the regulation is an activation or repression. When the database does not provide this information, if the regulator of the interaction is a TF we assume that the regulation is an activation, and if the regulator is a miRNA we assume that it is a repression. In the current version of BioNetUCR this network can be exported directly to Cytoscape for visualization (Figure S3, step 1).

##### BioNetUCR (step 2): Network Simplifications and Sensor Loops

BioNetUCR includes several commands for network simplification that we have used to reduce the complexity of our networks and enrich them for important interactions. The command ADD EDGE and DELETE EDGE enable to include or exclude particular interactions. The command DELETE NODE enables to delete specific nodes (e.g. species with poor fitting and no control on the genes of interest). Other commands implemented for reducing complexity of our networks are: DELETE AUTOREGULATED EDGES that removes arcs from one node to itself, DELETE SINK NODES removes nodes that are only sinks, DELETE SOURCE NODES removes nodes that are only sources and no other node regulates them. In addition, we implemented an algorithm of network construction to enrich the network with ‘Sensor Loops’ to favor the search of any kind of feedback or feed-forward interactions with a potential role in the network back regulation for the genes of interest such as those expected in the case of gene dosage compensation. A ‘Sensor Loop’ is a group of interactions starting from a gene of interest and returning to the same node after a determined number of arcs. To enrich the network, we implemented the following algorithm (pseudocode shown) where the inputs are the list of regulated genes of interest (*RegulatedGenes)* and *I,* which is the list of interactions (putative or experimentally validated) among species (*I*) and where every interaction is seen as an edge in a network, with a *source* node and a *target* node. The *OutputNetwork* is the list of edges of the output network, which is initially empty.

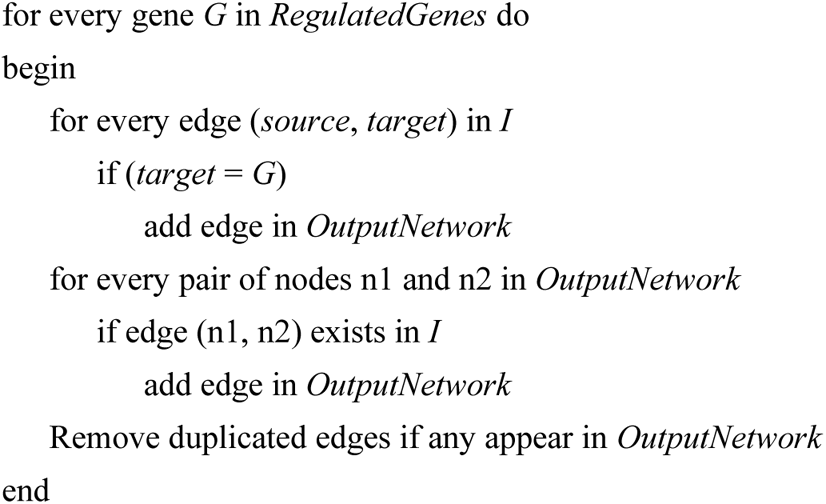

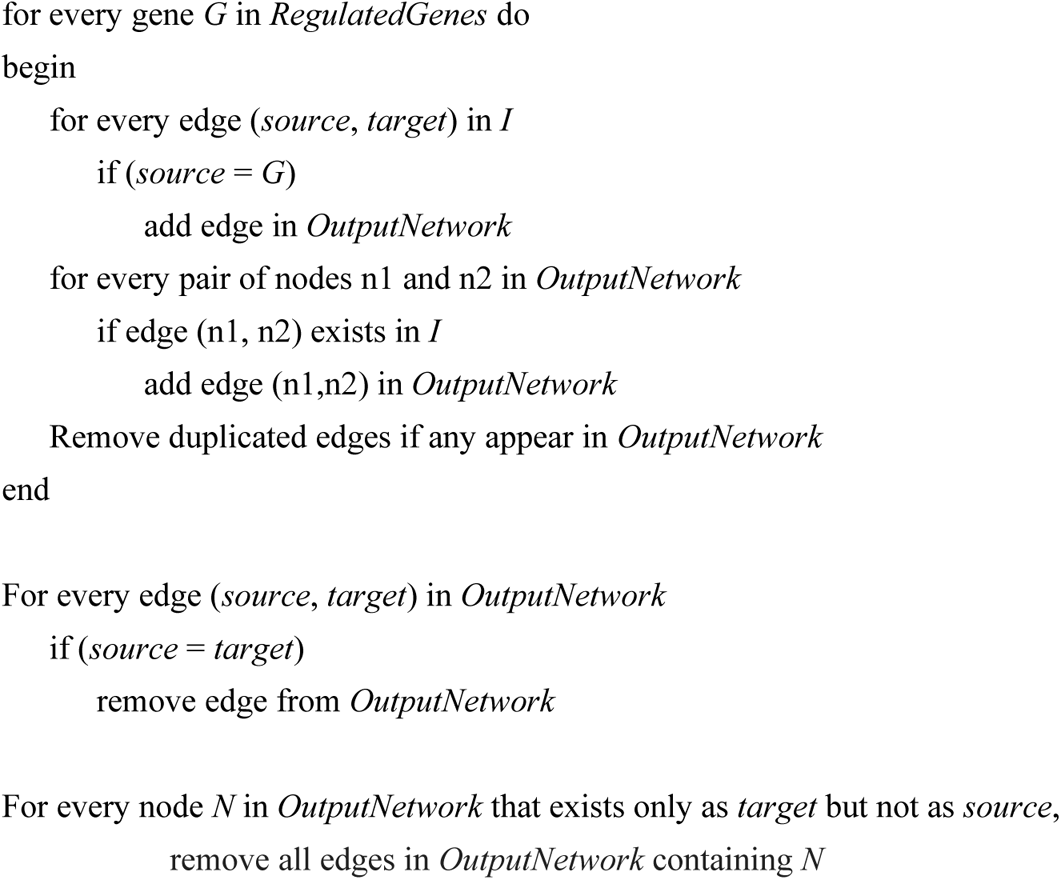

##### BioNetUCR (step 3): Conversion of network to ODE model

Since the gene dosage compensation phenomenon is gene expression related, we are interested in constructing a kinetic model from the graph. We now construct a system of ODEs from the graph representation of the regulation network. A script was written to automatically construct an SBML mathematical model. The network file (tab separated file of interactions) is input to the platform for model construction. Also, a script was written to create all the required reactions for all the species (i.e. nodes, either mRNA or miRNA), and write out a COPASI file. For each species of the graph, we write an equation computing the concentration or mRNA expression of this node. The list of all these equations defines a system of ODEs modeling the metabolic behavior of the regulation network. The schematic representations of these rate laws are shown in (Figure 2B). The rate of the transcription reactions includes all the effects of all positive and negative regulators according to a rate law of the form:

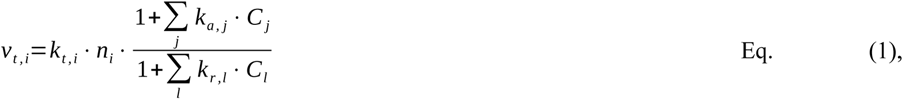

where *k_t,i_* is a basal transcription rate constant, *n_i_* is the copy number of gene *i*; the summations are over all species *j* that are positive effectors (numerator), and all species *l* that are negative effectors (denominator), each with a corresponding strength (*k_a,j_* or *k_r,l_*). Each species is also degraded with a basal first-order term, in the form:

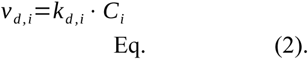

The model then consists of one ODE for each species *i*:

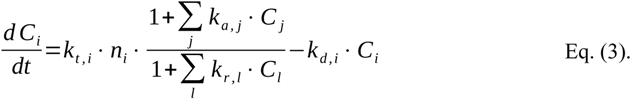

*Specific rate laws*

We have six different types of rate laws, one for each type of species: candidate gene, TF and miRNA.

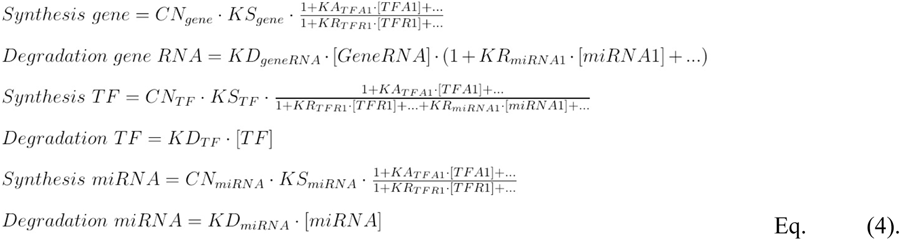

From the three types of equations defined, we detect three different types of elements: copy numbers (CN), rates (K) and concentrations ([TFA1], [GeneRNA], [miRNA]). There is a copy number parameter for each species of the graph. There are four different types of rates o parameters to be fitted against experimental data: synthesis (KS), degradation (KD), activation (KA) and repression rates (KR).

In order to model the total regulatory effect, we define the following parameters to be estimated later against experimental data. For a TF we define an activation rate (KA_TF_) that represents the rate at which the TF activates o intensifies the expression of the TF’s targets. Likewise, we define a repression rate (KR_TF_) representing the rate at which the TF suppresses or silences its targets when the TF has a repressing role. For a miRNA we also define a repression rate since miRNAs are assumed to only repress its targets (KR_miRNA_). The single regulatory effect of one TF or one miRNA over all of its targets is the product of the concentration of the TF or miRNA multiplied by the corresponding rate. Given that the concentration of a regulator, the activation rate and the repression rate are all zero or positive, the single effect of one regulator is also zero or positive. Following this definition, we have only one repression rate for a miRNA, therefore a given miRNA is assumed to exert the same repression rate over each of its targets.

The concentrations of the regulators that are part of the total regulation effect factor of the equation play an important role in the modeling of the regulatory effects. For an activator TF, as its concentration increases, so increases the activation effect and the concentration of the TF’s regulated gene. In the same way, as the concentration of a miRNA increases, so increases the repression effect causing a decline in the concentration of the miRNA’s regulated gene, either by slowing down its synthesis or by accelerating its degradation. The concentrations also play an important role in the modeling of common network motifs such as feedback loops, feedforward loops or other types of loops. In a loop like A->B->C->A, the concentration of A is part of the regulatory factor of B’s equation, the concentration of B is part of C’s equation, and C’s concentration is part of A’s equation. The interplay of these three equations models the dynamics of the loop in the graph.

For a candidate gene, the mRNA concentration equation has a synthesis part and a degradation part. The synthesis equation of the candidate gene is the product of three factors: a) the gene copy number, b) the synthesis rate of the gene, a parameter to be fitted by the Parameter Estimation task, and c) the total regulatory effect exercised by TFs on the candidate gene which in the graph is represented by all the incoming arcs of the candidate gene where the arc’s regulator is a TF. The total regulatory effect factor is usually >1 since most TFs are activators but it could be <1 when the regulating TFs are mainly suppressors. The degradation equation of a candidate gene is also the product of three factors: a) the mRNA concentration of the gene, b) the degradation rate of the gene, another parameter to be estimated, and c) the total regulatory effect of miRNAs on the gene. Because the degradation equation is a product and the total repression effect is one of the factors, the value of the repression effect is usually >1 since miRNAs are suppressors, thus exercising an accelerating effect on the degradation.

The mRNA expression equations for a TF or miRNA are slightly different from the equation of a candidate gene. These two equations also have a synthesis part and a degradation part but instead of having the repressing effect exercising an accelerator effect in the degradation part, they have the repressing effect in the synthesis part applying a decelerating effect in the synthesis part. The degradation equation for a TF or miRNA is the product of only two factors: a) the mRNA concentration of the TF or miRNA, and b) the degradation rate of TF or miRNA. The synthesis equation for a TF or miRNA is the product of three factors: a) the copy number of the TF or miRNA, b) the synthesis rate of the TF or miRNA, and c) the total regulatory effect exercised by others TFs and miRNAs on this TF or miRNA which in the graph is represented by all the incoming arcs of this TF or miRNA where the arc’s regulator is a TF or miRNA.

We modeled the total repression effect of miRNAs in the degradation equation of a candidate gene as the sum of 1 plus the single effect of each of the miRNAs repressing that gene. This value is always greater or equal than 1. When there are no miRNAs repressing the gene, this factor would be 1. These values model the expected function of the total repression effect in the degradation equation by accelerating gene degradation.

We defined the total regulation effect of TFs in the synthesis equation of a candidate gene as a ratio between the total activation effect of the TFs divided by the total repression effect of the TFs. The total activation effect is the sum of 1 plus the single effect of each of the TFs activating the gene. Likewise, the total repression effect is the sum of 1 plus the single effect of each TFs repressing the gene. When there are no TFs regulating the gene, this ratio is 1. When there are TFs activating or repressing the gene, if the total activation effect is greater than the total repression effect then the ratio is greater than 1, hence accelerating the synthesis of the gene. If there are more repression effect than activation effect, the ratio is less than 1 slowing down the synthesis. If both effects are equal, the ratio is 1 exercising no effect on the synthesis. These ratio’s values model the expected behavior of the interplay between the activating and repressing TFs in the synthesis equation of a gene in steady state.

While the total activation effect depends only on activating TFs, now the total repressing effect involves not only repressing TFs but also repressing miRNAs. Whether there are more total activating effect or more total repressing effect, the total regulatory effect would be >1 or <1, or exactly 1 if both effects are the same.

Having defined the equations of the species and identified the different elements of these equations, translating the graph of the regulatory network into an SMBL model (Hucka et al., 2003) is straightforward. An SBML model follows the XML standard (Hucka et al., 2003) (W3C/XML, 2008) and have several sections for the different elements of the SBML model.

##### BioNetUCR (step 4): Generation of experimental file

The SBML model is imported into COPASI, an application for analyzing biochemical networks like the miRNA-TF regulatory network. The platform also contains a script to extract NCI60, CCLE or TCGA data (breast cancer in the current version of BioNetUCR). For our NCI60 models the platform adds 59 sets of experimental data to this model, one set for each cell line of NCI60 coded as individual experiments to fit steady-state simulations named Experiments.tab. The odd row numbers contain the headers including a first set of columns for the copy number of each species and a second set of columns for the expression data of each species. The pair row number include the data for each of the cell lines of the corresponding data set.

##### BioNetUCR (step 5): Parameter estimation

The platform creates all the required reactions for all the species (i.e. nodes, either mRNA or miRNA), and write out a COPASI file that includes the experimental data ready for estimating the parameters of the ODE model. The platform then adds the experimental data file and the corresponding information in the COPASI file for the parameter estimation task. All the dynamic parameters K described above are included as the fitting parameters and the data file is parsed into individual steady-state experiments where the columns corresponding to Copy Numbers are included as the corresponding independent parameters in the model and the columns corresponding to Expression Data are included both as the corresponding independent initial concentrations of the species and the dependent variables of transient concentrations of the corresponding species at steady state.

Next the platform invokes COPASI to open the resulting file for us to carry out parameter estimation such that the model reproduces the provided experimental data. The Hooke-Jeeves minimization algorithm in COPASI produces good results, although the recently incorporated NL2SOL algorithm works even faster. The estimation was actually carried out in parallel in a cluster using Condor-COPASI (Kent et al., 2012) for the large networks and in normal PCs for the minimal models.

##### BioNetUCR (steps 6-7): Removal of non-fitted/weak regulators

To characterize the resulting model we designed a strategy combining Sensitivity Analysis and the evaluation of goodness-of-fit. A general sensitivity analysis for all the parameters is included on the model variables (species concentrations) and we extract the total control of each species on the regulated genes of interest (target genes) (step 6). In addition, we assess the goodness-of-fit of the RNA expression of the model to the RNA expression of the experimental data. To do that, we extract the ‘parameter estimation result’ report and imported in a functionality of BioNetUCR to parse the file, extract the experimental and simulated data comparison and perform a *t*-test for paired samples with α=0.05 to compare the model’s RNA expression of the 60 cell lines of a species with the corresponding experimental RNA expression. The pairing is on the cell lines. By plotting the total control (sensitivity analysis) against the p-values of the species it is possible to identify the species with the worse fitting and the lowest control as possible candidates to be removed from the model. We systematically removed one-by-one (step 7) the node with the lowest control on the genes of interest and the lowest p-value (first miR-106a and then miR145 for the NCI60 network) in BioNetUCR to generate a new network and COPASI model. A new Parameter Estimation task is run for the modified model until the difference between the experimental and the simulated data is not significant (T-student, p > 0.05). The presumption is that the experimental data do not explain the discarded species and they therefore do not belong to the model.

##### Assessment of gene dosage compensation

Once a fitted model is obtained, using the COPASI model we can manipulate the copy numbers of our target genes using the parameter scan function. To evaluate if a gene is compensated in the fitted model, we run a *Parameter Scan*, varying the copy number of the gene from 1 to 5 and computing the respective RNA expression. A non-compensated gene has a near linear fold increase in RNA expression level as its corresponding copy number increases linearly. We consider a gene compensated if the increase in RNA expression follows a sublinear increase as the copy number increases linearly. This experiment was repeated under different conditions (see results) to determine the most important species controlling the genes of interest and obtain a minimal model of gene dosage compensation.

##### Model of *MYC/STAT3* compensation in CCLE

The CCLE data was obtained from https://depmap.org/portal/download/. The CNV data included gene level copy number data, log2 transformed with a pseudo count of 1 and HGNC and Entrez ID identifiers. We obtained the RNAseq gene expression data for protein coding genes, also Log2 transformed with a pseudocount of1 and the same identifiers (“DepMap, Broad (2021): DepMap 21Q3 Public. figshare. Dataset,” n.d.). The miRNA expression data was also obtained from Depmap (Cancer Cell Line Encyclopedia Consortium, 2015; J et al., 2012). We also obtained the CCLE ABSOLUTE copy number analysis results of 2019 (Ghandi et al., 2019) and mapped the CNV data of the corresponding miRNAs to the raw segmented copy-number profiles in the .seg file format (hg19) of CCLE DNA Copy Number using the GRCh38.87 version of the genome. A full interception of all datasets was performed to have a complete dataset for 904 cell lines and 18431 instances including 541 miRNAs. The modeling approach was very similar to the one used with NCI60 but was performed at 3 different levels: pan-CCLE, tumor-type specific and cell line specific by delimiting the corresponding data sets.

##### Calculation of Compensation Index

We implemented a script using the COPASI API (*Application Programming Interface*, http://copasi.org) and its bindings for C# of .NET platform (https://dotnet.microsoft.com) to perform the fitting to individual experiments as a tool to characterize the model heterogeneity. We used a COPASI model (.cps) as basis and the experimental file (tab separated text file) with the rows containing the experimental data for each cell line (CCLE) or patient (TCGA). The fitting is executed with the Parameter Estimation task on each row corresponding to one experiment, choosing previously the NL2Sol optimization algorithm. The fitted values for all species and each experiment are tabulated in an Output CSV (*Comma-separated values*) file. This CSV file is input for the Parameter Scan task to perform individual parameter scans using the Parameter Scan task of COPASI by increasing the gene copy number from 1 to 5 and obtain a second output file for the simulated values. The respective “gene expressions” *in silico* were captured and normalized for each case (TCGA) or cell line (CCLE) dividing each of them by the *in silico* expression at a copy number of 1. The compensation index was calculated by dividing by 5 the normalized expression of each model at copy number of 5. A compensation index close to 1 indicates very weak compensation and a compensation index close to 0 indicates a strong compensation *in silico*.

##### Minimal model of MYC/STAT3 for TCGA

For the TCGA data, we developed scripts in Python to access the Application Programming Interface (API) of the Genomic Data Portal (GDC) to extract the data files “Copy Number Segment”, “Gene Expression Quantification”, “miRNA Expression Quantification” and their corresponding metadata including the UUID (Universally Unique Identifier) of each case (patient) and biospecimen information (sample, portion, analyte, aliquot). Afterwards we generated a script to process the Copy Number Segment mapping to the GRCh38.87 to obtain the CNV data for each gene and miRNA. The gene expression data comes in Fragments per Kilobase of transcript per Million mapped reads (FPKM), which normalizes read count by dividing it by the gene length and the total number of reads mapped to protein-coding genes. The miRNA expression data comes in counts normalized to reads per million mapped reads (RPM). Finally, we extracted the data for the species of the minimal models of *MYC*, *STAT3*, *STAT5A* and *STAT5B* dosage compensation and created the COPASI data files integrating the data of cases and biospecimens (using their UUIDs) for copy number and expression for each gene and miRNA. We performed the single-experiment parameter-estimation (see above) to obtain a mathematical model for each experiment and performed individual parameter scans increasing *MYC* copy numbers from 1 to 5 to calculate the respective Compensation Index as stated above.

### SUPPLEMENTARY SECTION

#### A miRNA/TF interaction network connects all candidate genes but its structural analysis reveals that only those with transcription factor function can be subjected to gene dosage compensation

We performed a correlation analysis of the copy numbers of target genes with the expression levels of miRNAs and transcription factors in order to identify possible regulators responding to the dosage of our candidate genes. We calculated the correlation coefficients between the Z-scores of copy number variations of the candidate genes and the Z-scores of miRNA/TF expression data across the NCI60 panel. As depicted in Figure S2A, there are both miRNAs and TF with positive or negative correlation to the candidate copy numbers, suggesting that many miRNAs and TFs are potentially involved in the regulation of the candidate expression in response to their gene dosage.

We asked whether this potential regulation is exerted by direct or indirect interactions. To evaluate whether there are any reported or predicted connections between these candidate genes and miRNAs/TF, we searched for putative and validated interactions between our candidate genes and miRNA/transcription factors in several databases (See STAR Methods). After filtering the correlation matrix of Figure 1A by these reported interactions, we obtained the correlation coefficients for direct gene-miRNA and gene-TF interactions as shown in Figure 2SB. Within those remaining correlations, we found that the highest positive correlation with miRNAs is only 0.34 for [(*ZNF217* copy number) to (miR-26b expression)] and the highest anti-correlation with TFs is −0.63 for [(*STAT3* copy number) to (HOXB13 expression)] followed by −0.5 for [(*MYC* copy number) to (*STAT3* expression)]. These results suggest that most correlations and anti-correlations between copy number and miRNA/TF expression should be due to indirect interactions; it is unlikely that the direct interactions alone would be responsible for gene dosage compensation of their corresponding candidate genes given those fairly low correlations.

A mechanism of gene dosage compensation will require a way to “sense” the changes in gene copy number and a type of feedback to exert the compensation. Therefore, we asked if those many miRNAs and TFs may form regulatory feedback loops to potentially compensate the dosage of our candidate genes. First, we constructed a network of putative and validated regulatory interactions based on the information available on the miRNA-target and TF-target databases cited above. In addition, we included the interactions between miRNAs and TFs using the information available at Transmir (Tong et al., 2019) and CircuitsDB 2 (Friard et al., 2010b). This generated a network with 612 nodes and 47122 interactions, which connected all 21 candidate target genes with 428 TFs and 163 miRNAs (Figure S2B).

In order to examine the target/miRNA/TF network for the presence of regulatory interactions, we searched for motifs with systems-level properties including positive and negative feedback loops (between miRNAs and TFs), coherent feedforward loops and incoherent feedforward loops (Vera et al., 2013). To quantify these interactions, we wrote an algorithm to search for cycles (STAR Methods).

We identified a total number of 5,054 putative regulatory motifs. For example, miR-15A participates in a coherent feedforward loop inhibiting both SEMAD3 and BRCA1, a positive regulator of BRCA1. Also, BRCA1 forms an incoherent feedforward loop, activating the transcription of SEMAD3 but also miR195, a negative regulator of SEMAD3. In addition, miR15A forms a negative feedback loop with BRCA1 (Figure S2C). In total, this network includes 156 negative feedback loops between TFs and miRNAs, 2337 coherent feedforward loops and 2561 incoherent feedforward loops formed by the interaction of TFs and miRNAs with the target genes. The analysis of this network topology indicates that several putative regulatory loops link all the candidate target genes.

The regulatory motifs with potential systems-level properties to mediate dosage compensation are widely present within this putative network. However, from the construction of this network topology, it is easy to infer that this network could sense changes in copy number of target genes only if they also have a TF-function (red colored interactions, Figure S2C). Otherwise, the remaining target genes are dead ends of the network, having only input arcs but no outputs. Among the 21 candidate target genes, only 7 of them correspond to transcription factors: *MYC*, *FOXC1*, *FOXF2*, *STAT3*, *STAT5A*, *STAT5B* and *ZNF217*. This structural analysis of the network topology indicates that for the remaining genes, the current network of interactions does not include a direct regulatory loop able to explain a potential dosage compensation. However, the gene dosage compensation of candidates with TF function might also impact the expression levels of other genes.

For the candidate genes with TF function, the identification of many potential regulatory loops offers a putative explanation of gene dosage compensation. Indeed, *MYC*, *ZNF*217 and *STAT3* are the most connected genes in the network, indicating that they participate in multiple regulatory loops. These results showed that a putative network of target-miRNA-TF interactions connects all the compensated genes but only those with TF function might be able to trigger a signal and get a compensatory response from that network. Due to the complexity of this regulatory network, a quantitative systems-level approach is required to identify regulatory loops mediating gene dosage compensation.

